# How attention saves energy in vision

**DOI:** 10.64898/2026.03.18.710397

**Authors:** Eivinas Butkus, Zhuofan Ying, Nikolaus Kriegeskorte

**Affiliations:** Department of Psychology, Columbia University, New York, NY, USA; Department of Neuroscience, Columbia University, New York, NY, USA; Department of Electrical Engineering, Columbia University, New York, NY, USA; Zuckerman Mind Brain Behavior Institute; Columbia University, New York, NY, USA; NSF AI Institute for Artificial and Natural Intelligence

## Abstract

Attention has long been thought to enable efficient vision,^1–8^ yet it requires additional neural machinery and energy. Whether attention yields net energetic benefits—after accounting for the cost of control—has never been demonstrated. Here we show that attentional control can substantially improve whole-system energy efficiency in a model of primate visual processing. Our model, EAN (“Energy-efficient Attention Network”), implements attention as recurrent top-down multiplicative gain over features, space, and time. EAN is optimized using a joint objective combining task performance and neurobiologically grounded energy costs accounting for action potentials and synaptic transmission across all components,^9–11^ including the attentional control circuitry itself. On a visual-category-search task requiring joint identification and localization of a target, EAN learns to focus its energy dynamically on task-relevant locations and features, reducing total energy use by up to 50% at matched accuracy and enabling flexible trial-by-trial trading of accuracy against energy. The model variant combining feature-based and spatial attention is most efficient and best captures human errors and difficulty judgments. EAN generalizes to classical attention tasks, replicating canonical effects of attention on firing rates, variability, and noise correlations,^12^ and patterns of V4-to-V1 feedback suppression.^13^ Our work connects a cognitive function (attention), a neural mechanism (gain modulation), and a neurobiological constraint (metabolic cost) in a single mechanistic model that explains how selection and recurrence enable flexible, energy-efficient vision.

---

Psychologists have long proposed that visual attention enables efficient use of limited neural resources.^1–8^ Attention is thought to select relevant aspects of the sensory evidence for prioritized processing in a cognitive bottleneck,^5,14^ thus reducing the energetic cost of vision by processing only a subset of the sensory information deeply.^6^

However, how exactly attention saves energy has remained a mystery. Attention requires additional machinery (a controller, top-down connectivity) and additional energy for running these components. The notion that energy is saved by selection also raises the question of what the selection is based on. If a feedforward pass already computes all the visual features,^15–18^ why recompute a subset of them?

This puzzle matters because vision is energetically expensive. The human brain uses about 20W, approximately one-fifth of the body’s energy budget.^19^ The cortex expends most energy for synaptic transmission, action potentials, and the maintenance of resting potentials.^9–11^ Vision is particularly energetically expensive in primates because their visual system occupies a large portion of cortex.^20^ Animal brains are thought to be highly optimized for energetic efficiency.^21–25^ To understand biological vision, therefore, we must consider energetic costs.

Previous work has primarily focused on how variables are efficiently *represented* in the brain, leaving out the costs of *computing* those variables from raw sensory signals. For instance, efficient neural coding principles^26–32^ explain how neurons maximize information per unit metabolic cost of the population code. Sparse coding is a canonical example demonstrating how primary visual cortex maximizes stimulus information per spike.^29^ However, efficient coding principles address only the final representational state, neglecting the substantial costs of computing that representation through hierarchical processing across multiple brain regions and iterative refinement over time. The principles of *efficient neural computation*—how brains optimize for the full cost of inference and adapt to changing energetic constraints and task demands—remain poorly understood.^33,34^

Attention is a key mechanism that likely plays a major role in efficient neural computation. William James influentially described attention as “the taking possession by the mind, in clear and vivid form, of one out of what seem several simultaneously possible objects or trains of thought”.^1^ The mind, thus, *chooses* content and computations. In modern terms, attention can be defined as the set of internal control mechanisms that actively select which aspects of the sensory input are processed and what computations are performed on them, given the goals and resources of the organism. In the context of vision, attention can select features, spatial locations, moments in time, or objects—giving rise to established forms of visual attention: “feature-based”,^35^ “spatial”,^36^ “temporal”,^37^ and “object-based”.^38^

Systems neuroscientists have found that attention modulates neural activity in primate visual cortex.^13,14,39–44^ Cellular neuroscientists have identified mechanisms—balanced synaptic input, shunting inhibition, and other gain control processes^45–49^—that could implement attentional selection.^50,51^ Beyond binary selection, neural signals can be continuously modulated to engage a graded trade-off, where higher-gain signals afford greater precision at greater metabolic cost. This is possible because gain modulation mechanisms can preserve the statistics of intrinsic noise,^45^ such that increasing the gain increases the signal-to-noise ratio.^52^ Brains can thus allocate more energy, and devote more spikes, to better represent important signals.

Yet how exactly attentional modulation signals are computed and how they save energy during visual tasks has remained a puzzle. To understand how attention saves energy in vision, we need to connect insights from cognition and neuroscience in a unified mechanistic model. The model must be able to perform a meaningful visual task and demonstrate that attention yields efficiency gains, while accounting for the full metabolic costs of computation.

Accounting for the full costs of inference requires moving beyond the efficient-coding paradigm. We introduce a general energy-accounting framework that measures action potentials and synaptic transmission across all components and time steps of a task-performing neural network. Unlike previous proxies for energy use,^53,54^ the synaptic transmission costs we introduce operate at the level of individual synapses, capturing activity-dependent costs that can be masked when excitatory and inhibitory inputs cancel. For these costs to be meaningful, we incorporate biologically plausible neural noise^55^ into all computations. Without noise, a neural network model could trivially minimize energy by scaling down all signals without affecting performance. However, such a model would not capture biological reality and would fail if implemented in neuromorphic hardware, where noise is an essential part of the challenge. The presence of neural noise establishes a fundamental energy-accuracy trade-off, where larger weights yield larger activations, which cost more energy but have the benefit of higher signal-to-noise ratios. Our energy-accounting framework is general and can be applied to any neural network model—a step from efficient coding toward principles of efficient neural computation.

To demonstrate how attention can save energy in vision, we apply our energy-accounting framework to a novel neural-network model family we call EAN (“Energy-efficient Attention Network”). The key intuition is that a relatively cheap attentional control circuit can substantially improve whole-system energy efficiency by modulating the visual hierarchy of representations. EAN combines established components: a convolutional neural network (CNN) approximates the primate visual hierarchy,^15–18^ while a recurrent neural network (RNN) captures visual inference dynamics.^56–60^ The RNN also implements the “attentional controller”, which can modulate the visual hierarchy via top-down multiplicative gain signals.^61–63^ The top-down gain can target specific features, locations, or moments in time—integrating “feature-based”, “spatial”, and “temporal” attention within a single modular architecture. EAN is optimized under a joint objective that measures both task-performance errors and energy costs, and learns to dynamically allocate energy in a graded fashion to the features and locations of a particular image that matter for perceptual performance.

To see is “to know what is where by looking”.^64^ Handling the combinatorics of what and where efficiently is a core computational challenge for any visual system. To capture these essential elements of vision, we test EAN on a novel visual-category-search (VCS) task, in which the subject has to find a handwritten digit (target category) among distractor letters, with uncertainty about both the target’s class (“what”, 0-9) and location (“where”). Knowing where the target is would make the identification of the digit easy. Knowing what digit the target is, conversely, would simplify localization. Defining the target at the category level (“find the digit”) entails a dual uncertainty about what and where that is ubiquitous in everyday life. For instance, finding a good option at a buffet takes time because preferred foods can have many different appearances (uncertainty in what) and be located in many different places (uncertainty in where).

A naive algorithm for VCS that evaluates all identity hypotheses for all locations would be highly inefficient. EAN’s attention mechanisms instead efficiently eliminate locations and features, dynamically focusing energetically costly scrutiny on the task-relevant locations and features. This adaptive inference process substantially improves energy efficiency and enables flexible trial-by-trial accuracy-energy trade-offs, where the same model can spend more energy to achieve higher accuracy. We compare model variants and find that combined feature-based and spatial attention is most efficient and best explains human errors and difficulty judgments.

EAN’s attentional controller can be adapted to perform a diverse set of visual tasks. EAN generalizes to classical attention tasks and replicates canonical electrophysiological effects of attention on firing rate, variability (mean-matched Fano factor), and noise correlation.^12^ The model also captures how V1 firing rates are affected by optogenetic suppression of V4 feedback.^13^

We provide a mechanistic solution to a long-standing question—how attention improves the efficiency of vision—by connecting a cognitive function (attention), a neural mechanism (multiplicative gain control), and a neurobiological constraint (energy metabolism) in a single model. The model explains how attention—despite requiring additional machinery—can yield net energy savings by controlling the gain of features and locations in a hierarchy of visual representations. Beyond attention, the energy accounting framework introduced here provides a general approach to studying how neural circuits optimize the full cost of computation.

## Visual-category-search (VCS) task

In our VCS task, the subject had to find a handwritten digit among distractor letters and report its identity (“what”) and location (“where”) (Fig. 1a). Unlike classic visual search tasks with a known target,^65^ VCS requires overcoming dual uncertainty: subjects do not know either the identity (0-9) or the location of the target digit before the search image appears.

**Figure 1:**
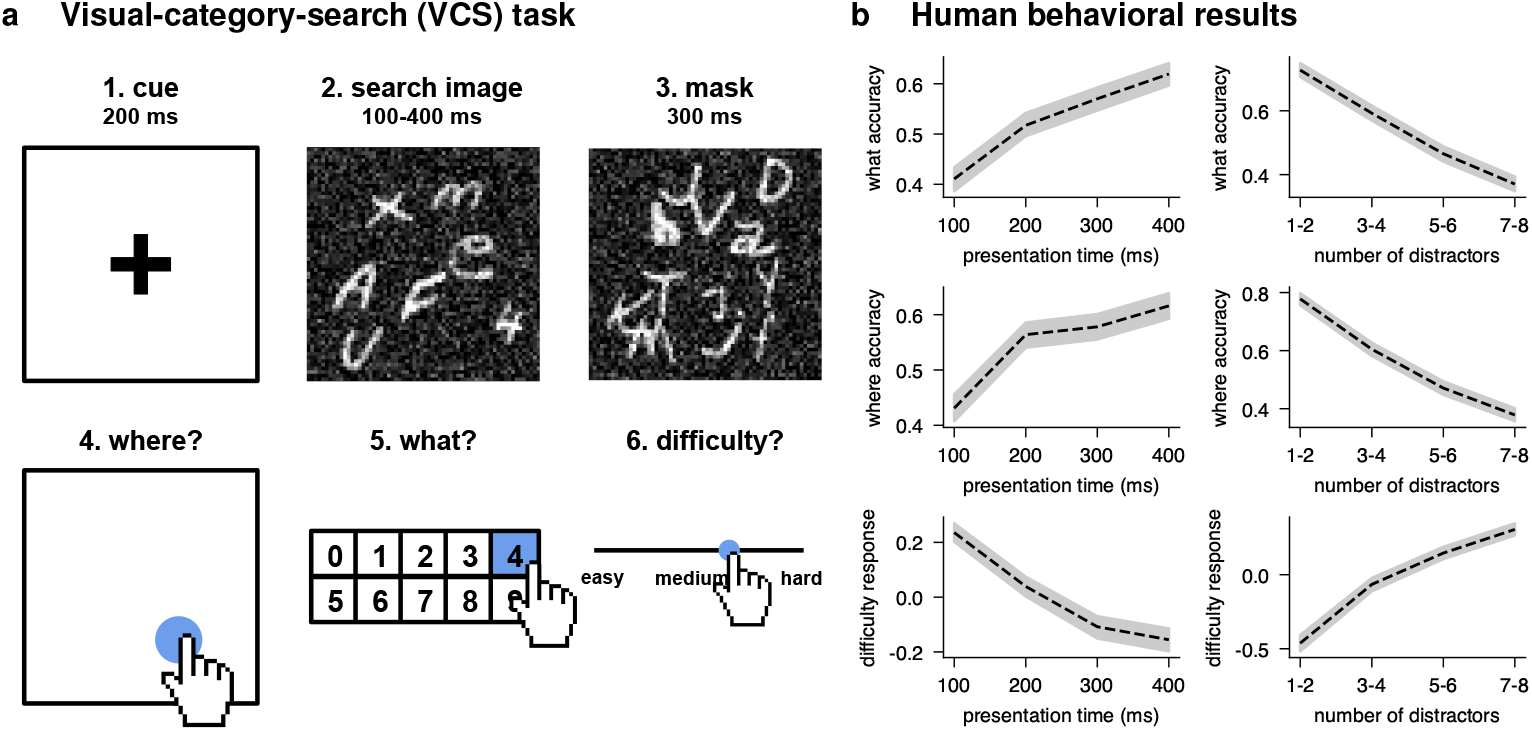
**a** The visual-category-search (VCS) task. After a brief presentation (100-400 ms) of the search image (with one target digit and 1-8 distractor letters), the subject has to report *where* they saw the digit, *what* class it belonged to, and the *difficulty* of that particular search. **b** Human behavioral results (N=18, 389 trials per subject) as a function of presentation time (left) and number of distractors (right). Accuracy in both what and where components of the task increases with longer presentation durations, and decreases with the number of distractors. Difficulty judgments (z-scored) show the reverse trend: trials with longer durations and fewer distractors are rated as easier. Shaded regions represent 95% confidence intervals across trials and subjects.

Human subjects viewed each search image for 100-400 ms, followed by a mask meant to disrupt iconic memory and terminate recurrent processing. They then reported the digit’s location (*where*) and identity (*what*) and rated the trial’s *difficulty*. Models received only the search image (no cue or mask) and predicted the digit’s class and location.

## Energy-efficient Attention Network (EAN)

Our model EAN (Fig. 2) tests the hypothesis that an energetically inexpensive attentional controller can substantially improve net energy efficiency of vision. All versions of the model share the same backbone consisting of a “visual hierarchy” (implemented as a three-layer CNN) and an “attentional controller” (implemented as a three-layer RNN). On each time step, the model receives a 64 × 64 image as input and processes it using the visual hierarchy followed by the attentional controller. There are two recurrent pathways in EAN: (a) lateral connections within the attentional controller and (b) top-down connections that modulate the visual hierarchy via multiplicative gain. At every time step, we read out the class of the digit (“what”) and its location (“where”) from the hidden state of the last layer of the attentional controller.

**Figure 2:**
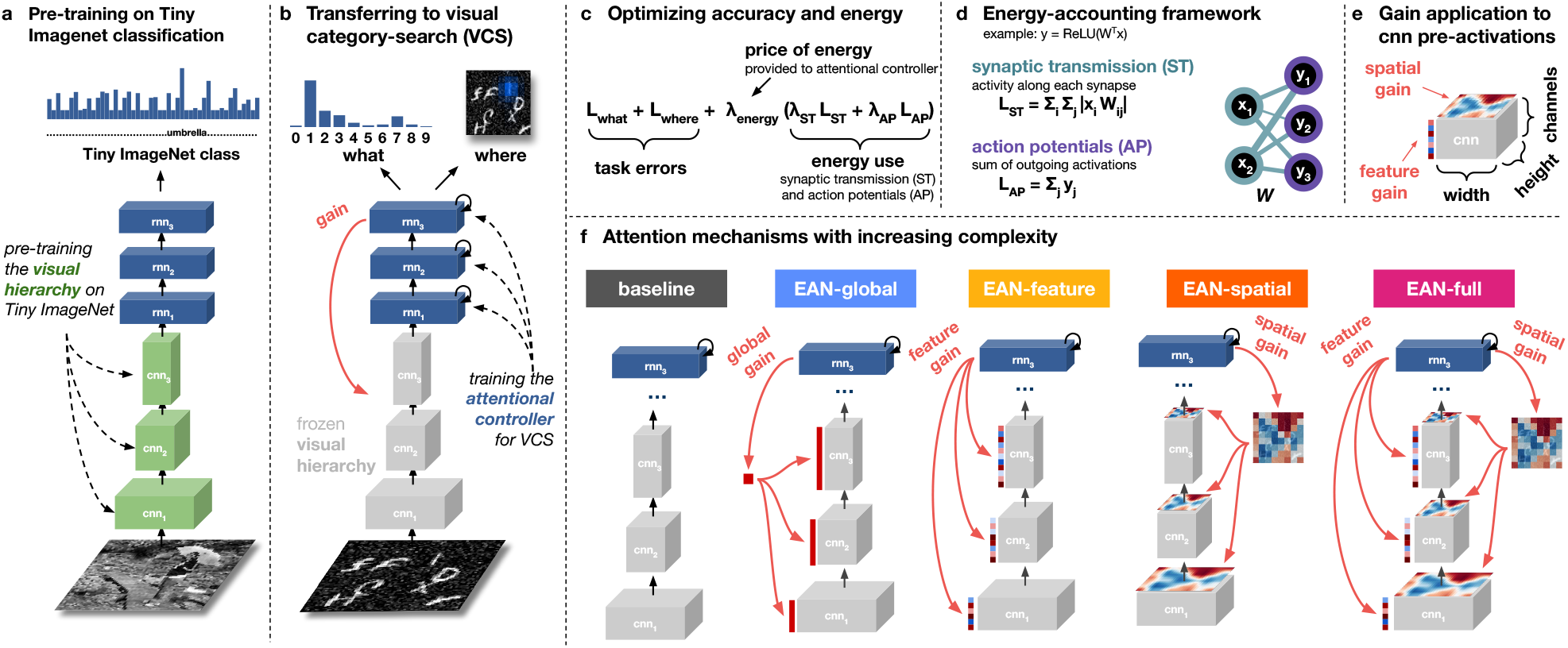
EAN (“Energy-efficient Attention Network”). **a** The visual hierarchy is pre-trained for object classification on TinyImagenet to obtain general-purpose visual features. **b** Convolutional weights in the visual hierarchy are frozen (pre-attentional selectivity is fixed) and only the attentional controller, gain mechanisms and readout are trained for VCS. **c** We optimize a joint cost of task errors and energy use based on neurobiologically plausible measures of synaptic transmission and action potentials, where *λ*_energy_ modulates the relative cost of energy. **d** Energy-accounting framework. Energy used for synaptic transmission depends on the activations of the previous layer (pre-synaptic firing rates) and the weights (synaptic strengths), while energy for action potentials depends on the activations (post-synaptic firing rates). **e** Multiplicative gain is applied to the convolutional pre-activation tensor. Feature gain is multiplied along the channel dimension, while spatial gain is multiplied across width and height. **f** Different versions of EAN implement increasingly more sophisticated attention mechanisms, capturing classical cognitive notions of “feature-based”, “spatial” and “temporal” attention within the same modular architecture.

Biological visual systems learn visual representations that can be flexibly deployed for a diverse set of behavioral goals.^66^ To obtain general-purpose visual features, we pre-train the visual hierarchy within a feedforward (single time step) version of EAN for object classification on the Tiny Imagenet dataset (Fig. 2a). We then *freeze* the convolutional weights, fixing the pre-attentional selectivity of all units in the visual hierarchy. Finally, we train only the attentional controller, the gain mechanisms and the readouts on the VCS task, unrolling the model for four time steps (Fig. 2b).

## Energy-accounting framework

The optimization objective for EAN combines cross-entropy loss terms (as surrogate objectives for accuracy in the what and where tasks) and differentiable measures of energy consumption (Fig. 2c). The energy costs, grounded in neurobiology, account for action potentials and synaptic transmission across all model components and across all time steps.^9,11^ The energy-accounting framework introduced here is general and can thus be applied to any neural network.

For action potentials, we treat post-rectified-linear-unit (ReLU) activations as proportional to neural firing rates and minimize their sum over space, time, and inputs, reflecting the total metabolic cost of generating spikes (Fig. 2d). This is consistent with efficient coding models and standard sparseness penalties on the activity of the units.^29^ For synaptic transmission, the appropriate cost metric is less obvious. Previous proxies for synaptic transmission cost—L1 penalties on weights^53^ or absolute pre-activations^54^—fail to capture activity-dependent costs at the level of individual synapses. We instead compute the sum across all synapses of the absolute products of pre-synaptic firing rates and synaptic weights (Fig. 2d). This activity-dependent measure of the energetic cost of synaptic transmission is summed over time points and inputs.

An artificial neural network without internal noise could arbitrarily scale down all its weights and activations to minimize the energy costs defined above without affecting performance. In a neural network simulated on a digital computer, the only limit to down-scaling of activations is floating-point precision. In contrast, biological neural systems have multiple sources of internal *noise*,^55^ all of which corrupt neural signals. We therefore add normally distributed noise to all pre-activations in EAN, so that lower pre-activations yield worse signal-to-noise ratios. The noise establishes a biologically realistic trade-off between conserving energy (by making weights and activations smaller) and maintaining signal precision. After noise is added to the pre-activations, we apply ReLU non-linearity. Each ReLU activation is divisively normalized by the pooled activations within its neighborhood defined by a fixed Gaussian kernel^51^. The normalized activations correspond to the neuronal firing rates, whose energetic cost is measured by a term in the objective.

We combine task errors and energy use in a single loss function using the energy-cost factor *λ*_energy_ (Fig. 2c), which we also refer to as the “price of energy”. When *λ*_energy_ = 0, we recover the standard deep learning cross-entropy objective (minimizing task errors). When *λ*_energy_ *>* 0, the model minimizes energy use along with errors. Intermediate values of *λ*_energy_ encourage the model to balance accuracy and energy use. High settings of *λ*_energy_ compromise task performance. The price of energy can either be “fixed” across trials (log *λ*_energy_ ∈ {−12, …, −5}) or “flexibly” sampled on every trial (log *λ*_energy_ ∼ Uniform(−12, −6)), ranging from accuracydominant to energy-dominant regimes. To enable EAN to flexibly account for the variable cost of energy on a trial-by-trial basis, the value of *λ*_energy_ is provided as input to the attentional controller at the beginning of the trial. When trained in the “*λ*_energy_-flexible” regime with a distribution of energy prices, a single EAN instance can therefore use attentional gain modulation to dynamically trade off between accuracy and energy based on the trial-specific energy cost.

## Modular attention architecture

The attentional controller computes top-down gain signals that multiplicatively scale pre-activations in CNN blocks throughout the visual hierarchy (Fig. 2e). Inspired by neurophysiological findings that gain modulation preserves internal noise levels,^45^ the scaling is done *before* noise is applied, so gain typically leads to higher signal-to-noise ratio but also incurs a higher energetic penalty.

We implement increasingly sophisticated forms of attention within a modular architecture (Fig. 2f). The baseline model lacks top-down modulation entirely and has a single gain parameter that scales pre-activations independent of current inputs or time step. EAN-global implements temporal attention through a single dynamically computed scalar gain value that uniformly modulates all units at each time step. EAN-feature adds feature-based attention, enabling the controller to prioritize specific convolutional filters at each time step.^62^ The gain for each filter is then applied uniformly across the visual field.^67^ Instead of feature-based attention, EAN-spatial employs spatial attention, boosting or suppressing all convolutional filters at specific locations. Finally, EAN-full combines feature and spatial gain, enabling the most flexible attentional control. The attentional controller outputs a spatial gain map and a feature gain map, and the gain applied to each unit is the product of its spatial gain and its feature gain. The modular design allows us to isolate and compare the contributions of each attention mechanism to both accuracy and energy efficiency.

We hypothesized that EAN-full, combining both feature-based and spatial attention, would achieve the best energy-accuracy trade-offs because the product of the two attentional filters can eliminate a large proportion of the less relevant units, enabling highly selective choice of the most relevant computations. Fig. 3 demonstrates EAN-full inference dynamics across four time steps, given different energy costs *λ*_energy_. For an intermediate energy cost, the model initially attends broadly to gather information, then dynamically focuses on task-relevant locations and features to efficiently resolve uncertainty about what and where.

**Figure 3:**
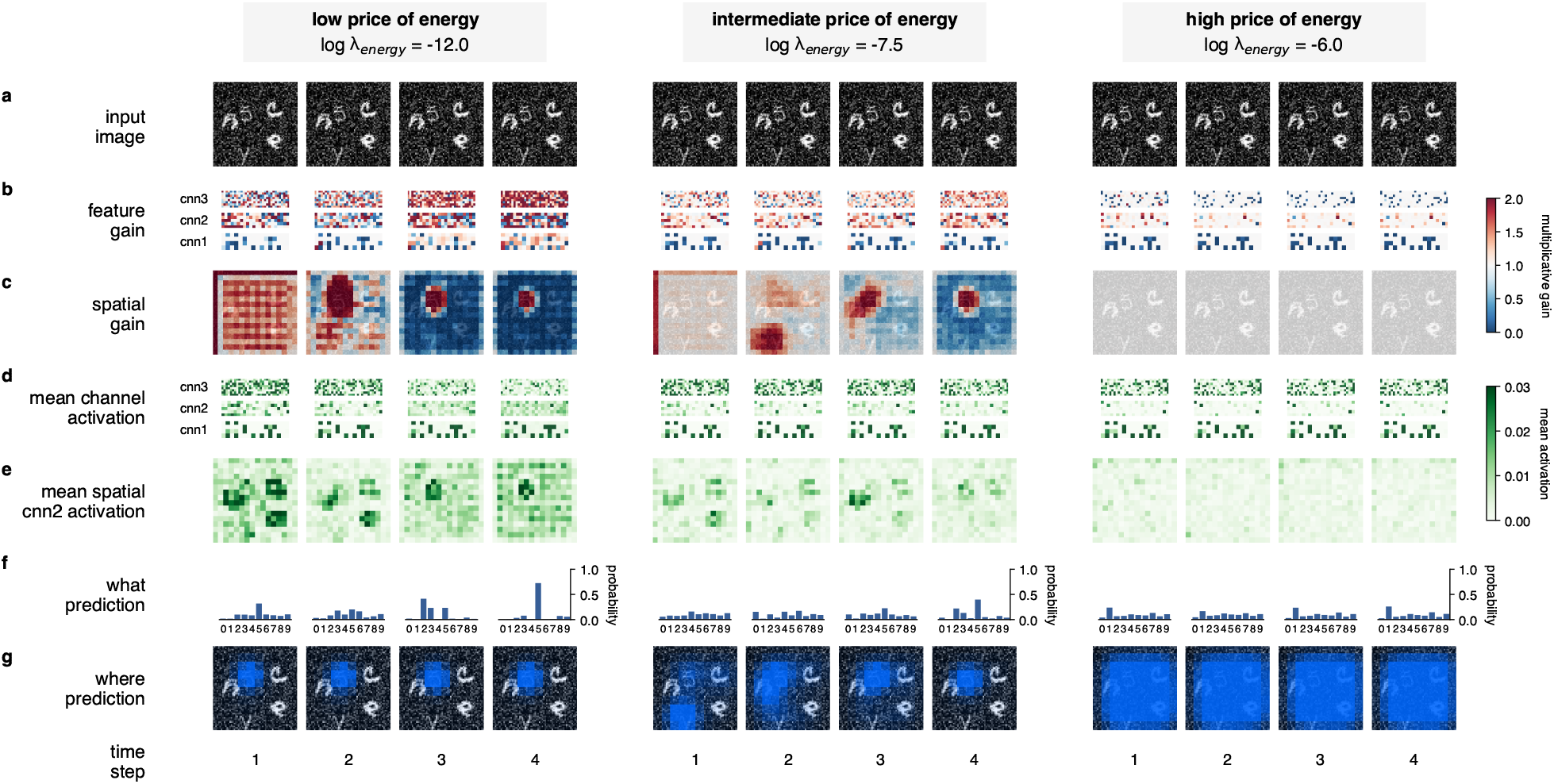
EAN-full inference dynamics in VCS under the *λ*_energy_-flexible regime, where a single model instance is trained to handle a distribution of energy prices. We plot model behavior for low (left), intermediate (center), and high (right) price of energy *λ*_energy_. **a** Input image (repeated across time step in VCS). In this particular trial, the target is digit 5. **b** Feature gain for each convolutional layer. Patterns are relatively sparse and inhibitory (especially for cnn_1_), which may explain why feature gain brings about large energy savings (Fig. 4). **c** Spatial gain. The model eliminates implausible locations and focuses on the target digit, while inhibiting distracting letters. **d** Mean channel activation for each convolutional layer. **e** Mean spatial activation for cnn_2_ (averaged across channels). Since we treat activations as firing rates, darker green corresponds to higher energy use. **f** Model predictions for the target digit class (what). **g** Model predictions for the target location (where). The three columns illustrate how EAN can flexibly modulate its own activity and trade accuracy for energy. When energy is cheap (left), EAN uses higher activations and arrives at a confident, correct prediction. At an intermediate energy price (center), EAN balances energy use with task performance. When price of energy is high (right), EAN suppresses its activity, sacrificing task performance.

## Attention improves energy efficiency

We first assess how attention affects energy efficiency when models are trained with *fixed* energy-cost factor *λ*_energy_ (Fig. 4a). In this regime, each model instance is optimized for a single constant price of energy (log *λ*_energy_ ∈ {−12, −11, …, −5}).

**Figure 4:**
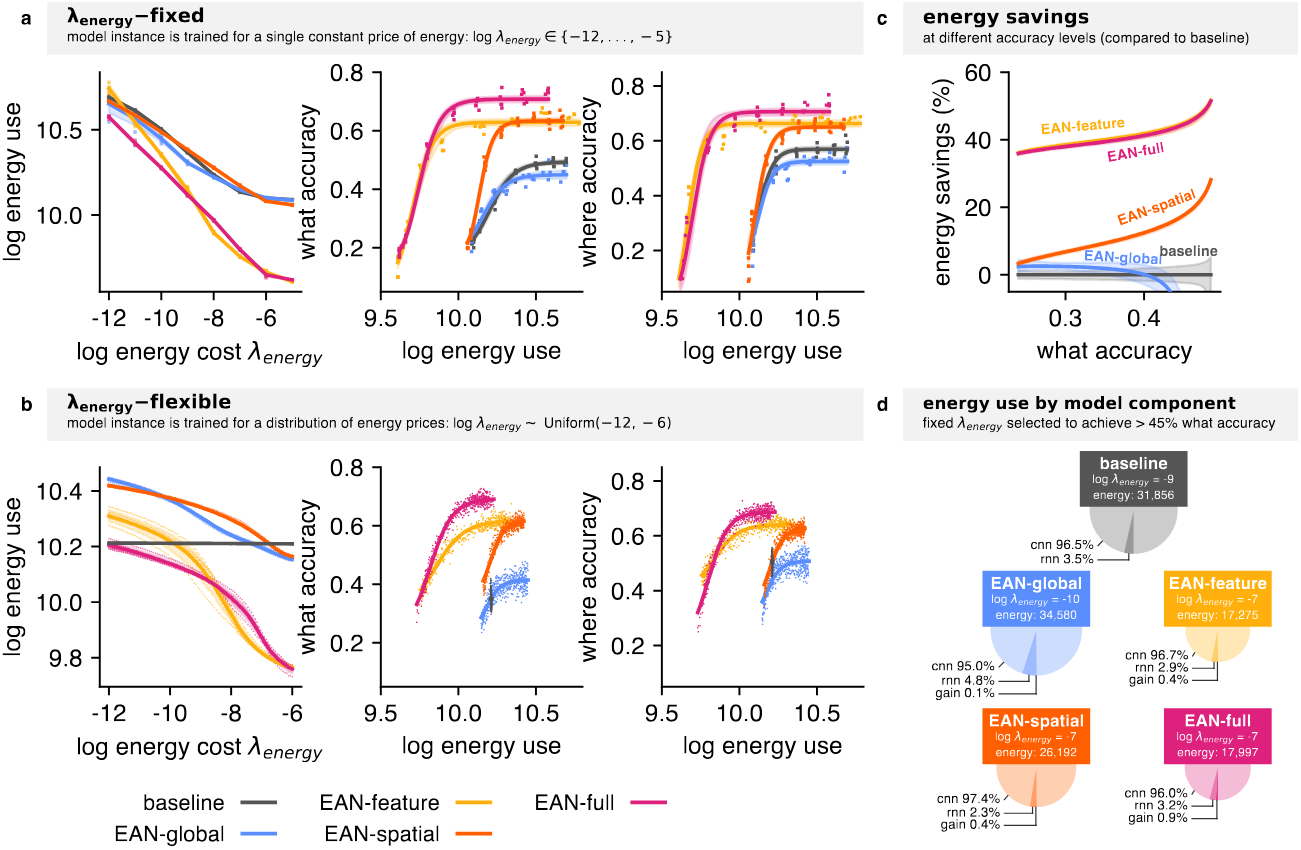
**a** Attention mechanisms improve energy efficiency. *Left*: Models trained with higher *λ*_energy_ (stronger energy penalty) use less energy (measured by action potentials and synaptic transmission). *Middle and right*: Higher energy use is associated with better performance in both what and where components of the task. Each point represents the average accuracy-energy profile for a model instance trained with a fixed price of energy (log *λ*_energy_ ∈ {−12, …, − 5)}. EAN versions with spatial and feature gain achieve superior energy-accuracy trade-offs compared to baseline (gray). Lines with shaded regions show logistic fits with 95% bootstrapped confidence intervals across trials. **b** Attention enables flexible energy-accuracy trade-offs. When trained with a distribution of energy prices (log *λ*_energy_ ∈ Uniform(− 12, − 6)), a *single* EAN instance can dynamically adjust its energy use based on the cost specified at inference time. In contrast, the baseline model without attention must commit to a single operating point. **c** Attention mechanisms (particularly feature-based gain included in EAN-feature and EAN-full) yield up to 50% net energy savings compared to the baseline model (from the “fixed” *λ*_energy_ regime). **d** Energy use breakdown by component for models trained in the “fixed” regime (*λ*_energy_ selected to achieve *>* 45% what accuracy). Components are: “cnn” (convolutional layers in the visual hierarchy), “rnn” (recurrent layers in the attentional controller), “gain” (top-down gain computation). Area of the pie is proportional to total energy use by the model. Relatively cheap attention mechanisms (RNN and gain computation) enable large reductions in net energy use.

As expected, higher energy prices lead to lower energy use (Fig. 4a, left). Importantly, model instances that use more energy achieve higher accuracy in both what and where components of the VCS task (Fig. 4a, middle and right). Our optimization approach yields models that span the full spectrum from energy-dominant to accuracy-dominant solutions.

Consistent with previous work,^61^ attention mechanisms improve *task performance* (Fig. 4a, middle and right). At high energy use, where models can fully leverage their computational capacity, models with feature-based attention (EAN-feature) and models with spatial attention (EAN-spatial) achieve substantially higher accuracy than the baseline model. EAN-full, which combines both types of attention mechanisms, achieves the highest accuracy for what and where judgments—higher than either attention mechanism alone—suggesting that spatial and feature-based attention provide complementary benefits. Interestingly, EAN-global (the model that implements temporal attention only) does not improve performance over the baseline model. This suggests that for static stimuli, temporal modulation alone (uniform gain across features and locations) provides little benefit over baseline.

Critically, attention improves *energy efficiency*. Models with attention achieve better energy-accuracy trade-offs (Fig. 4a, middle and right). The energy-accuracy frontier improves systematically with the specificity of attentional control: selective gain over features or locations outperforms the no-gain baseline and uniform gain (EAN-global), while combined feature-and-spatial control achieves the best trade-off.

The energy savings from attention are substantial (Fig. 4c). Feature-based attention (included in both EAN-feature and EAN-full) is associated with the largest energy savings, achieving the same accuracy as baseline while using up to 50% less energy. Crucially, computing attentional control signals requires only a small fraction of total energy (Fig. 4d). This reflects the high cost of processing many features at every location across multiple layers of the visual hierarchy.

EAN explains how attention mechanisms can improve energy efficiency. The key insight is that the “attentional controller” can be a relatively small and energetically cheap circuit yet bring substantial gains in efficiency.

## Attention enables flexible use of energy

We next trained models with energy-cost factors sampled trial-by-trial rather than fixed (log *λ*_energy_ ∼ Uniform(− 12, − 6)). The attentional controller is informed about the price of energy on each trial: it receives *λ*_energy_ as input. Models with attention mechanisms can therefore modulate their activity according to the trial’s price of energy. In contrast, the baseline model without attention must commit to a single energy-accuracy operating point, finding a compromise that works for the distribution of energy prices.

Fig. 3 qualitatively demonstrates the adaptability of a single EAN-full instance trained in the *λ*_energy_-flexible regime. When energy is cheap (log *λ*_energy_ = − 12), the model uses higher gain magnitude and reaches confident predictions quickly. When energy is expensive (log *λ*_energy_ = − 6), the same model instance utilizes attention to inhibit visual hierarchy activity, sacrificing task performance. At intermediate energy prices (log *λ*_energy_ = − 7.5), the model deploys moderate gain signals and dynamically explores hypotheses about what and where while balancing accuracy and energy—arguably the most biologically plausible regime.

As in the *λ*_energy_-fixed regime, EAN-full achieves the best energy-accuracy trade-offs (Fig. 4b, middle and right). However, here a *single* trained instance of EAN spans the full range of energy-accuracy trade-offs by adapting to trial-specific costs. One might expect that a model trained for a particular price of energy will have better performance at that price of energy than a *λ*_energy_-flexible model. In fact, the *λ*_energy_-flexible variant of EAN-full achieves roughly equal accuracy at each price. The baseline, lacking attentional control, converges to an energy-accuracy regime that represents a compromise across the distribution of energy prices (Fig. 4b).

Beyond the efficiency gains shown under fixed energy costs, attention can also enable dynamic, trial-by-trial adaptation to the changing relative costs of energy and accuracy. This flexibility is relevant for biological vision, where metabolic availability and task priorities change frequently.

## EAN captures human behavior

EAN captures multiple aspects of human behavior in the VCS task (Fig. 5). Below, we focus on models trained in the *λ*_energy_-flexible regime.

**Figure 5:**
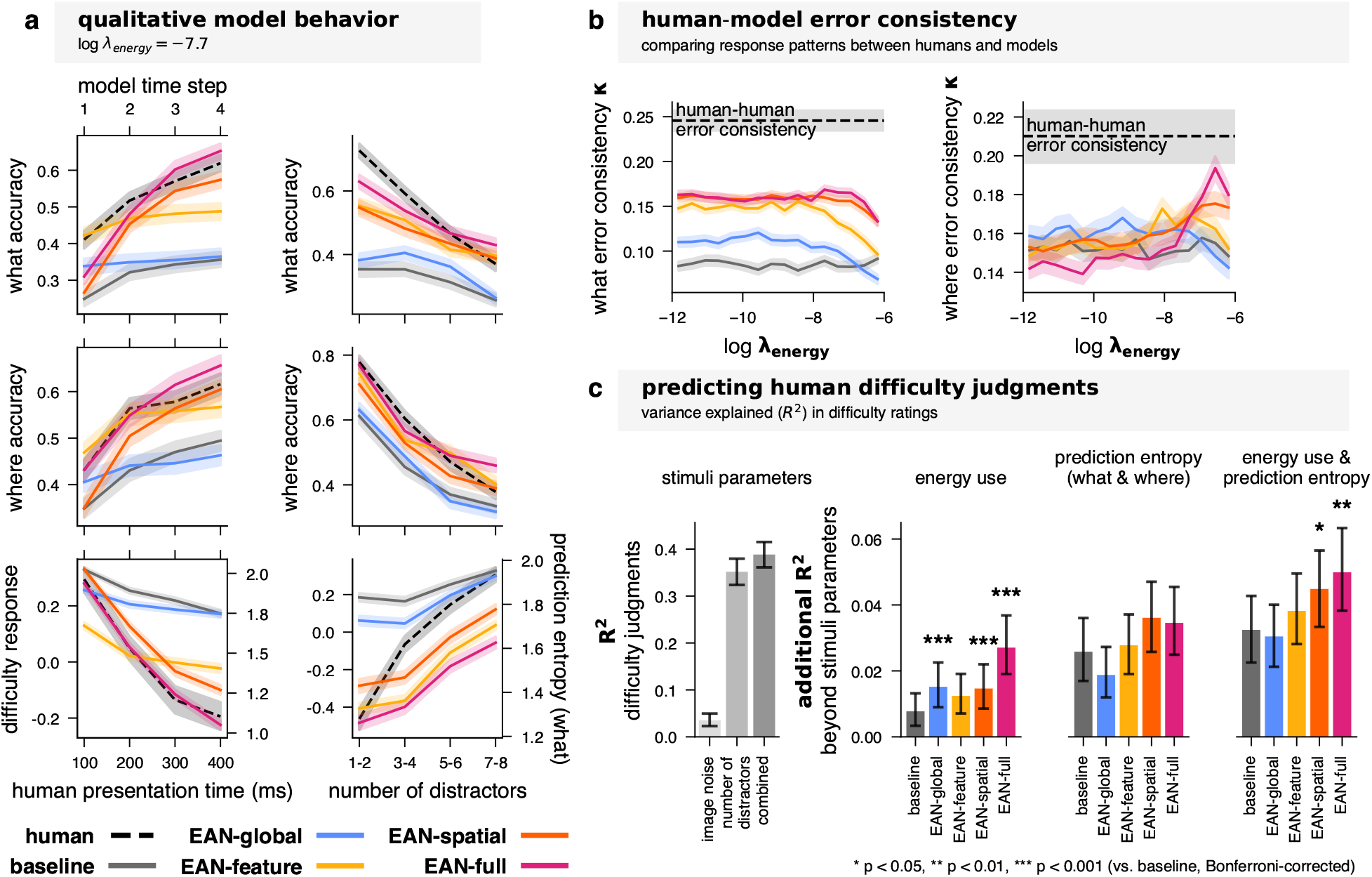
EAN-full, combining feature-based and spatial attention, best captures human errors and difficulty judgments. **a** Qualitative human-model comparison. Model behavior is plotted as a function of time step (left column) and number of distractors (right column) at an intermediate energy price (log *λ*_energy_ = − 7.7). Model prediction entropy in the what dimension (bottom row) serves as a qualitative proxy for human difficulty judgments. **b** What and where error consistency measured by Cohen’s kappa between humans and models, plotted as a function of energy price *λ*_energy_. Dashed lines with shaded regions indicate human-human error consistency with 95% confidence intervals. **c** Left: Variance in trial-level difficulty judgments explained by the amount of noise and the number of distractors in the search image. Right: *Additional* variance in difficulty judgments explained by model energy use and prediction entropy (model uncertainty). Significance stars indicate comparisons against baseline (Bonferroni-corrected).

First, we compare human and model behavior *qualitatively* (Fig. 5a), where all models operate under an intermediate price of energy (log *λ*_energy_ = − 7.7). We plot model behavior as a function of time step (left column) and as a function of number of distractors (right column). We use model prediction entropy in the what dimension as a qualitative proxy for human difficulty judgments (bottom). All model versions capture broad trends in human behavior. However, spatial attention (used in EAN-spatial and EAN-full) seems critical in capturing the dynamics of visual inference across time. Similar to humans, models with spatial attention use time as a resource to arrive at a better answer,^68^ while other models (baseline, EAN-global, EAN-feature) reach peak performance within approximately two time steps.

We used Cohen’s kappa to measure *error consistency* in the what and where components of the task.^69,70^ Error consistency measures trial-by-trial agreement between two classifiers while controlling for agreement expected by chance, given their respective accuracies. We plot error consistency as a function of energy price *λ*_energy_ (Figure 5b). Attention mechanisms improve error consistency, especially in the what component of the task. EAN-full achieves the highest error consistency, peaking at *κ* = 0.17 for what error consistency (at log *λ*_energy_ = −7.7, humans *κ* = 0.25) and *κ* = 0.19 for where error consistency (at log *λ*_energy_ = −6.6, humans *κ* = 0.21).

Finally, we turn to human difficulty judgments. Prior work has shown that metabolic activity scales with visual processing demands,^71^ and we hypothesized that subjective difficulty judgments might partially reflect trial-specific energetic costs alongside other factors such as the number of distractors and overall uncertainty. First, stimulus parameters alone (amount of noise and number of distractors) explain 39% of the variance (*R*^2^ = 0.39). We then tested whether model-derived measures—trial-level energy use and prediction entropy (quantifying model uncertainty in what and where tasks)—could explain *additional* variance in difficulty judgments. For each model variant, we selected the price of energy *λ*_energy_ that maximized the additional variance explained for each set of covariates used (see Extended Data Fig. 2 for results across the *λ*_energy_ range).

All models explained significant additional variance beyond stimulus parameters. For energy use, EAN-full showed the largest improvement (Δ*R*^2^ = 0.027), significantly outperforming baseline (Δ*R*^2^ = 0.008, *p <* 0.001). For prediction entropy, EAN-spatial (Δ*R*^2^ = 0.036) and EAN-full (Δ*R*^2^ = 0.035) both captured substantial additional variance. However, differences from baseline (Δ*R*^2^ = 0.026) were not significant after correction for multiple comparisons. The combination of both energy use and prediction entropy showed the strongest performance overall, with EAN-full (Δ*R*^2^ = 0.050) and EAN-spatial (Δ*R*^2^ = 0.045) significantly exceeding baseline (Δ*R*^2^ = 0.032, *p <* 0.01 and *p <* 0.05, respectively). These results suggest that human difficulty judgments reflect not only basic stimulus properties but also trial-specific internal processing demands, which are better captured by models with selective attentional mechanisms.

Together, these analyses demonstrate that EAN-full, combining feature-based and spatial attention, best captures the pattern of human errors and subjective difficulty judgments in VCS.

## EAN captures electrophysiology of attention

Beyond energy efficiency and human behavior, EAN replicates several electrophysiological effects of attention: canonical effects on firing rates, Fano factor, and noise correlation from Cohen & Maunsell [12], and effects on V1 firing rates following optogenetic V4 feedback suppression from Debes & Dragoi [13]. We implemented simplified four-time-step versions of the classical attention tasks used in the two papers. As for the VCS task, we kept the parameters of the pre-trained visual hierarchy fixed and trained only EAN’s attentional controller and gain mechanisms under the *λ*_energy_-flexible regime.

In Cohen & Maunsell [12], the monkey is presented with Gabor stimuli on both sides and must saccade to the stimulus that changes orientation, given an 80% valid attentional cue indicating which side is likely to change (Fig. 6a, top). For the model, instead of the dual *what* and *where* readout used in VCS, we implemented a “saccade” readout (“left”, “center”, “right”). EAN was trained to output “center” on all but the change frame, and “left” or “right” on the change frame (depending on which side changes).

**Figure 6:**
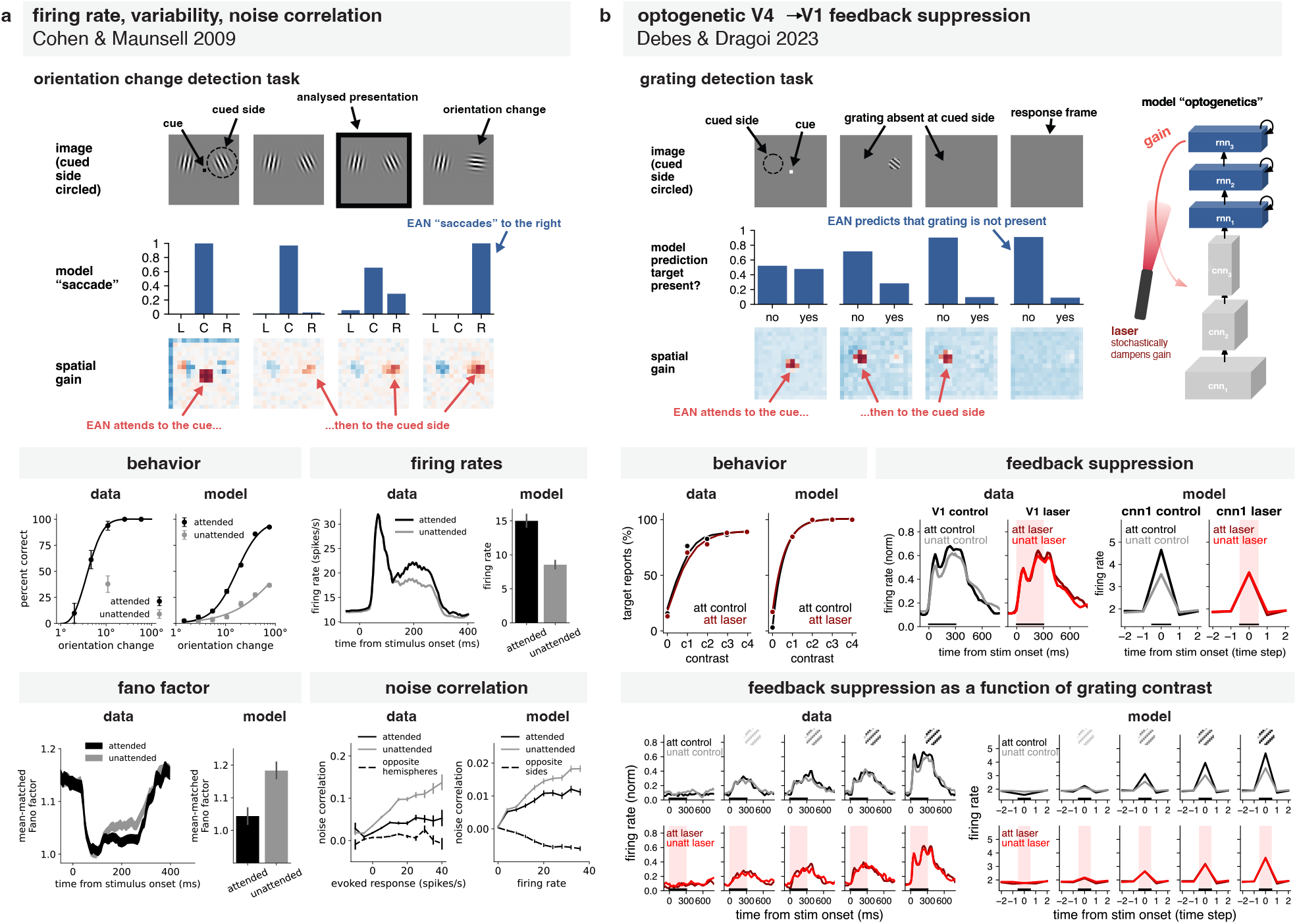
EAN generalizes to classical attention tasks, replicating canonical electrophysiological effects. Top: simplified versions of the tasks^12,13^—model input, output, and spatial gain across time steps. Bottom: data-model comparisons. Figures adapted from respective papers. **a** Replicating effects from Cohen & Maunsell [12]. The task is to saccade to the Gabor stimulus that changes orientation (attentional cue valid 80%). Model learns to perform the task by paying attention to the cue, then to the cued location. EAN replicates four canonical effects: higher accuracy for valid cues, increased firing rates at attended locations, decreased mean-matched Fano factor, and decreased noise correlation with attention. **b** Replicating effects from Debes & Dragoi [13]. The task is to detect whether a grating is present at the cued side (attentional cue valid 100%). On 50% of the trials, V4 to V1 feedback is suppressed using optogenetics on stimulus onset. We implement an analogous model “optogenetics” procedure, where we stochastically dampen the gain signal. EAN learns to perform the grating detection task by attending to the cued side. EAN replicates the main patterns: target reports increase with contrast, feedback suppression abolishes attentional modulation in V1, and the suppression effect increases with stimulus contrast.

EAN learned to perform the orientation-change detection task, exhibiting an interpretable spatial attention pattern: it first attends to the cue and then shifts attention to the cued location. Critically, EAN replicates four canonical effects of attention from Cohen & Maunsell [12] (Fig. 6a, bottom): (1) accuracy in the task is higher when the cue is valid, (2) firing rates are higher for units with receptive fields in the attended region, (3) mean-matched Fano factor decreases with attention, and (4) noise correlation—trial-to-trial variability shared between neurons when they respond to the same stimulus—decreases with attention.

In Debes & Dragoi [13], the monkey had to detect the presence of a Gabor-like grating of various contrasts on the cued side (Fig. 6b, top). The cue was 100% valid and the monkey had to completely ignore the uncued side. The authors showed that optogenetic suppression of feedback from V4 to V1 abolishes attentional modulation. We implemented an analogous feedback suppression procedure in EAN that stochastically dampens the gain signal from the attentional controller to the visual hierarchy. We trained the model with a simple binary readout to output “yes” if a target is present on the cued side, and “no” otherwise.

We replicate the main patterns of feedback suppression from Debes & Dragoi [13] (Fig. 6b, bottom): (1) target reports increase with contrast (but feedback suppression does not affect behavioral performance), (2) suppressing feedback abolishes attentional modulation in V1, and (3) the effect of feedback suppression increases as a function of stimulus contrast.

EAN is a brain-computational model that explains how attention can save energy in vision despite requiring additional components and energy. The model links the cognitive function of vision to its neurobiological implementation, using a biologically established gain control mechanism and explaining canonical electrophysiological signatures of attention and the effects of interventions using optogenetics.

## Discussion

We started with a fundamental puzzle: How can attention save energy if attentional control requires additional neural components and metabolic expenditure? Confirming a long-standing hypothesis that attention enables efficient vision,^6^ we provided a rigorous demonstration that a relatively cheap attentional controller can yield substantial net energy savings for the visual system as a whole. We defined neurobiologically grounded differentiable objectives for energy— accounting for both action potentials and synaptic transmission throughout all components. By optimizing a neural network model with a joint objective that balances task performance and energetic demands, we establish that attention can enable an efficient adaptive inference process, selecting relevant sensory signals for scrutiny and dynamically trading perceptual precision against energy consumption.

EAN’s architecture and efficiency objective give rise to a model that explains results of both human psychophysics and animal electrophysiological experiments. In the visual-category-search task, attention clearly improves energy efficiency. EAN achieves higher accuracy at lower energy costs than models without selective gain modulation (Fig. 4a). The energy savings are large: attention mechanisms can cut the energy use in half compared to the no attention baseline (Fig. 4c, d). The bulk of these savings is due to feature-based attention, while spatial attention provides complementary performance gains by progressively narrowing in on the most promising locations. Attention additionally enables a single model instance to dynamically trade accuracy for energy depending on current metabolic and task demands (Fig. 3, 4b). The model with combined feature and spatial attention (EAN-full) best captures human error patterns and subjective difficulty judgments in the VCS task (Fig. 5). EAN generalizes to classical attention tasks and replicates canonical electrophysiological effects of attention on firing rates, variability, and noise correlation, as well as recent findings from optogenetic suppression of V4-to-V1 feedback (Fig. 6). Together, these results position EAN as a mechanistic bridge between cognitive function, neural implementation, and metabolic constraints.

EAN generates new testable predictions about the attentional system in relation to metabolic costs. As a consequence of its modular attention architecture, EAN predicts that the metabolic savings afforded by feature-based attention are larger than those afforded by spatial attention (Fig. 4). This asymmetry has not been systematically explored in neuroscience. EAN’s prediction could be tested in macaques by selectively suppressing feedback^13^ from ventral prearcuate region (VPA), which is preferentially associated with feature-based attention,^72^ and frontal eye fields (FEF), which is preferentially associated with spatial attention.^73^ When task accuracy is matched, suppressing feature-based attentional feedback should result in greater increases in visual cortex metabolic expenditure (which should be reflected in a larger overall BOLD fMRI response) compared to suppressing spatial attentional feedback.

EAN also explains how attentional control enables biological visual systems to dynamically adjust how much energy they spend on a task (Fig. 4b). When task demands or metabolic constraints change, attention can down-regulate neural activity to save energy, accepting lower perceptual accuracy in return. This trade-off should emerge most clearly in visual tasks requiring joint feature and spatial selection such as VCS, where selective gain modulation affords the greatest range of accuracy-energy operating points. The prediction could be tested by combining VCS with fMRI to simultaneously track behavioral performance and metabolic expenditure across a wide range of reward magnitudes.

Recent computational models have shown that humans flexibly adapt their internal representations and computations to task demands,^74,75^ suggesting a form of cognitive resource rationality.^76,77^ However, the mechanisms implementing such flexibility—and the nature of the computational resources themselves—have remained abstract. EAN bridges this gap: it grounds high-level notions of selective attention and resource limitations in concrete neural mechanisms (multiplicative gain) and neurobiologically measurable costs (action potentials and synaptic transmission), while remaining an image-computable task-performing network that can be readily trained on new visual tasks, as demonstrated by generalization to classical attention paradigms.

Beyond cognitive science and computational neuroscience, our energy-accounting framework and findings about attention have implications for energy-efficient AI. Modern AI systems have embraced a relational attention mechanism as a core computational primitive,^78^ yet their energy demands remain immense compared to brains. Neuromorphic hardware aims to close this gap by implementing neural computation in physical substrates that respect the principles underlying biological computation.^79^ Our energy-accounting framework—measuring action potentials and synaptic transmission across all components and timesteps—provides an optimization objective aligned with the cost structure of neuromorphic hardware, and could guide the development of networks designed for efficient neuromorphic deployment. Moreover, our findings that selective attention can halve energy use in vision and enable flexible energy-accuracy trade-offs suggest a general design principle: incorporating a cheap attention controller and feedback to modulate signals can yield substantial energy savings in neuromorphic systems.

While we account for the cost of computing the gain signal, we may underestimate the metabolic cost of gain application, which has not been rigorously measured in the primate brain. Our energy-accounting framework is straightforwardly extensible, however: Additional cost terms can be incorporated as empirical estimates become available. Although EAN relies on top-down modulatory signals from the attention controller, top-down and lateral connections within the visual hierarchy are absent. Such connectivity is present in the primate brain and could further shape energy-efficient representations, perhaps learning to predict and cancel activity as in predictive coding.^54^ Finally, EAN implements a simplified visual hierarchy and operates in a controlled task environment; extending the framework to more naturalistic scenes and richer behavioral paradigms remains an important direction.

The energy-accounting framework introduced here—measuring action potentials and synaptic transmission across all model components and across time—is neurobiologically grounded and general. It can be applied to investigate how other mechanisms, beyond attention, contribute to metabolic efficiency across different tasks and brain areas, and whether optimizing energy efficiency improves model-brain alignment. In the context of EAN, the optimization of energy efficiency resolves a long-standing puzzle, revealing how our selective attention, which James described over a century ago^1^, enables efficient neural computation.

## Methods

### Visual-category-search (VCS) task

The goal in VCS for both human subjects and models was to determine the identity (“what”) and location (“where”) of a handwritten digit (“target”, 0-9) among handwritten letters (“distractors”). Digits were sampled from the MNIST dataset^1^, while letters were sampled from the EMNIST dataset.^2^

#### Generative model of the stimuli

Our stimuli are noisy grayscale 64 × 64 images consisting of one target digit and a number of distractor letters (1-8 for the human behavioral experiment and model evaluation, and 2-8 for model training). We start by sampling a target MNIST digit image (28 × 28) with its class label (0-9) from the MNIST dataset. We create the larger image 64 × 64 (“canvas”) and fill it with zeros. We sample the number of distractors from a discrete uniform distribution (1-8), and sample distractor letter images from the EMNIST dataset (28 × 28), while excluding letters that are most confusable with digits (‘i’, ‘l’, ‘o’, ‘q’, ‘s’, ‘b’, ‘z’, ‘g’). We downsize all of the digit and letter images to 16 × 16 and uniformly sample a location for each small image so that: (1) minimum center-to-center distance of 10 pixels is maintained to other letter/digit images and (2) each image remains fully within the canvas. We place the small target and the distractor images by adding them to their sampled locations in the big 64 × 64 canvas image (so overlapping items can create composite pixel values). We then set all pixels below *µ*_target img_ to *µ*_target img_, establishing a uniform background that does not make the target pop out. Finally, we sample the standard deviation of the image noise *σ*_img noise_ ∼ Uniform(0, 0.2), sample the noise value 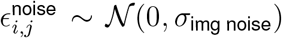 independently for each pixel *i, j*, add noise values to the image, and clamp image values to be within [0, 1].

#### Model training dataset

To train the model we sampled search images with their corresponding what and where targets from the generative model “on-the-fly” during training. We used “train” subsets of MNIST and EMNIST and each search image contained 2-8 distractors. For the what target, we created a one-hot target vector using the class of the MNIST digit from the MNIST dataset (0-9) and used cross-entropy as the loss function. For the where target, we created a soft 2D map by evaluating a 2D Gaussian kernel at 10 × 10 locations with the mean placed at the ground-truth location of the digit (the center where the 16 × 16 downscaled MNIST image was placed in the 64 × 64 canvas) and standard deviation *σ* = 1. We then normalized this heatmap to sum to one and used KL divergence as the loss function.

#### Behavioral experiment dataset

Behavioral experiment dataset (400 search images and 400 corresponding mask images) was used to evaluate both humans and trained models. We used “test” sets of MNIST and EMNIST to generate the behavioral experiment dataset. Each search image in the behavioral dataset contains 1-8 distractors. For each search image we also generated unique “mask” image, which was meant to disrupt recurrent visual processing and visual iconic memory in human subjects after predefined durations (100, 200, 300, 400ms) of the search image (Fig. 1a). We initialized the mask image canvas using the mean value of the search image. We then sampled 10-15 distractor letters (no digit), and used the same procedure of adding them to the canvas as for the search image. Finally, we used the same sampled noise standard deviation from the search image to generate the noise for the mask image.

#### Behavioral experiment

We implemented the behavioral experiment using the FlyingObjects package.^3^ The behavioral experiment consisted of 400 trials, and each subject saw the same set of images. We randomized the order of the trials for each participant. For each trial, we independently and uniformly sampled the presentation time from {100, 200, 300, 400} ms. The first 10 trials were designated as “training trials” with much longer presentation times (linearly going down from 2200 ms to 400 ms), which we excluded from the analysis. We also did not present the last trial due to an indexing issue, leaving 400 (all trials) − 10 (training) −1 (indexing issue) = 389 trials per participant for analysis.

We served the experiment to 20 subjects on Prolific and excluded data from 2 subjects that did not complete the full set of trials. The experiment was approved by Columbia University’s Institutional Review Board, and all participants provided informed consent. The median duration of the experiment was 45 minutes and the adjusted pay was $14.28/hr.

The structure of the trial was the following (Fig. 1a):

1. **Pre-trial**. 300ms empty screen before the trial starts.
2. **Cue**. A black cross in the middle of the screen presented for 200ms to allow subjects to anticipate the upcoming search image.
3. **Search image**. Presented for *t* ∼ 𝒰 ({100, 200, 300, 400})ms.
4. **Mask image**. Presented for 300ms. The mask was meant to “wipe” iconic memory of the search image and terminate recurrent processing.
5. **Where response**. Subjects had to click on an empty square to indicate their best guess where the target digit was. A blue bubble appeared to indicate where their click was registered. Note that during training trials, the bubble would appear green/red depending on whether the target digit was within 16 pixels of their click.
6. **What response**. Subjects had to click on one of the digit tiles. As with the where response, the tile would turn blue to indicate which digit class they selected. Note that during training trials, the digit tile would turn green/red depending on whether their selection was correct/incorrect.
7. **Difficulty response**. The subjects were also instructed to rate the difficulty of each trial on a continuous scale “Easy-Medium-Hard”. A blue bubble appeared after their click to indicate the continuous value of difficulty they selected.
8. **Post decision**. 200ms break before the next trial starts.

Our experiment was meant to study recurrent visual processing and the dynamics of covert attention of foveal vision within a single fixation. While we could not fully control the size of the stimuli since we served the experiment online, we estimate that the presented images extend around 2 - 3 degrees of visual angle (given normal monitor or laptop setup). Empirically, human accuracy in our experiment is relatively constant as a function of target eccentricity (Extended Data Fig. 1). This suggests that lower acuity of peripheral vision does not play an important or systematic role in our experiment.

Accuracy in the *where* component of the VCS task was computed by counting the proportion of trials in which the subject’s click location (or model’s simulated click location) was within a certain radius of the target. For humans, we set the max radius for a correct trial at *r* = 0.15, where the whole square image has an area of 1. For models, we simulated clicks by taking the argmax location of the 10 × 10 model prediction distribution and we used a smaller radius *r* = 0.07 to account for motor and memory noise in humans.

### Model architecture

All versions of the model share a common neural network architecture composed of a convolutional neural network (“visual hierarchy”), followed by a recurrent neural network (“attentional controller”). Besides the baseline model, all versions of the model also have 1-2 multi-layer perceptrons (“gain mechanisms”) that take the hidden state of the attentional controller as input. The output of the gain mechanisms is used to compose a multiplicative gain map that modulates the visual hierarchy.

#### Visual hierarchy

The visual hierarchy extracts general-purpose visual features from the image (the selectivity of the units is fixed after pretraining on TinyImagenet, see section on pre-training below) and passes those to the attentional controller.

The visual hierarchy is a convolutional neural network composed of three convolutional blocks (cnn1, cnn2, cnn3). The three convolutional layers within each block have 64, 128, and 256 feature maps (or channels) and kernel size 5, 3, 3 respectively. This results in the following activation tensor dimensionality (width x height x channels): input image (64×64×1), cnn1 (32×32×64), cnn2 (16×16×128), cnn3 (8×8×256).

These are the processing steps within one convolutional block:

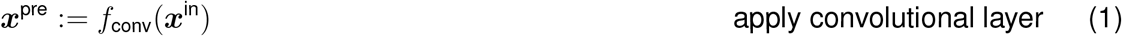

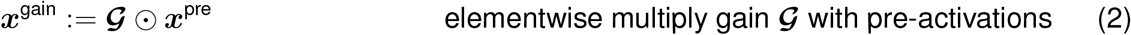

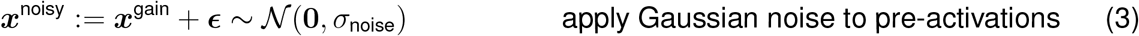

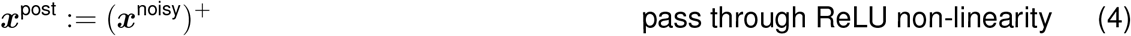

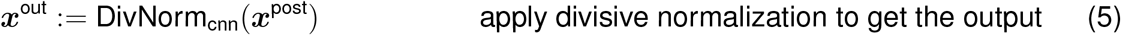

where divisive normalization^4,5^ for a CNN block is defined as:

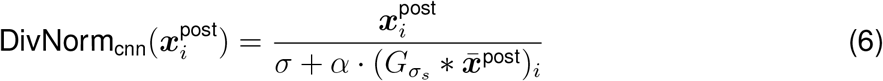

Here 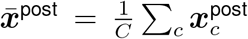 is the mean across channels, 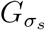 is a fixed spatial Gaussian kernel with standard deviation *σ*_*s*_ = 2.0 (pool size 5, weights sum to 1), *σ* = 1.0 is a semi-saturation constant, and *α* = 0.2 is a scaling constant. To ensure consistent normalization scales across layers, the pooled activation in the denominator 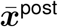 is divided by a running mean of channel-averaged activations, updated with momentum 0.1 during training. This ensures that, on average, the pooled term contributes a unit-scale value, so the denominator has a mean of approximately *σ* + *α* = 1.2 across all units.

#### Attentional controller

The attentional controller (1) integrates evidence from the visual hierarchy across time and (2) the hidden state of the attentional controller is provided as input to top-down gain mechanisms (excluding the baseline model without top-down modulation).

The attentional controller is a recurrent neural network composed of three recurrent blocks (rnn1, rnn2, rnn3). Each recurrent block learns how to update its hidden state (256-dimensional) across time, given incoming inputs from recurrent block below (or the last convolutional block for rnn1) and its own previous hidden state via a lateral hidden-to-hidden connection. Note that the output of the last convolutional block is flattened before being passed to the first recurrent block.

These are the processing steps within one recurrent block:

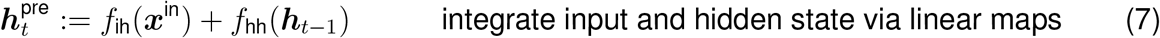

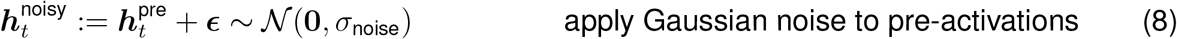

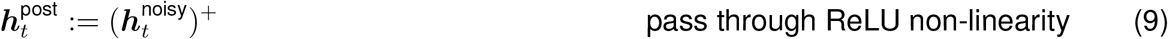

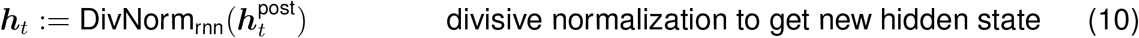

where divisive normalization for an RNN block is defined as:

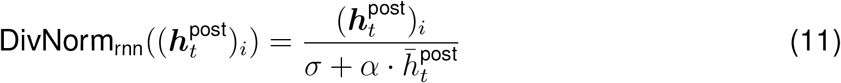

Here 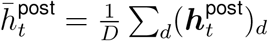 is the mean across the hidden dimension, with *σ* = 1.0 and *α* = 0.2 as for the CNN blocks. Unlike the CNN case, the RNN normalization pools globally across the hidden dimension rather than locally via a spatial kernel.

#### Gain mechanisms

Gain map ***G*** applied in convolutional blocks is computed using multi-layer perceptrons (MLPs, with one 256-dimensional hidden layer) that take as input the hidden state of the last recurrent block 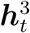.

The following equations describe how the gain maps are computed for different models:

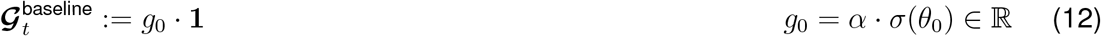

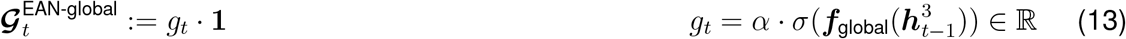

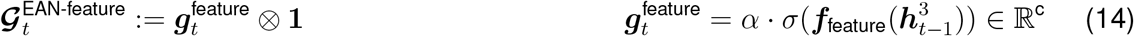

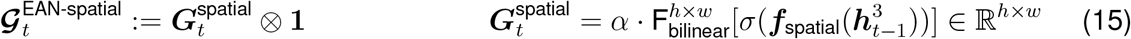

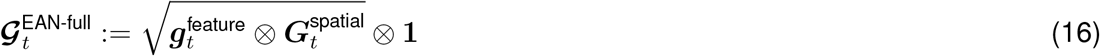

All ***f*** here are MLPs, 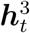 is hidden state of the final recurrent block for time step *t, σ* is the standard logistic function (sigmoid) applied element-wise, ⊗ is multiplication broadcasted across space or feature dimensions, **1** is a tensor of ones matching the dimensions of the convolutional layer pre-activation tensor ***x***_pre_, *α* = 2 is the scaling factor for the gain map. 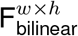 is bilinear interpolation that takes the 16 × 16 spatial gain map and scales it to *h* × *w* (the spatial dimension of the convolutional pre-activation tensor). The baseline model has a single gain parameter *θ*_0_ that can scale the pre-activations independent of input or time step.

We set *α* = 2, which scales the gain values to be in [0, 2] (from [0, 1] output of the standard logistic function). This *α* value ensures that identity gain operation (𝒢= **1**) is the default behavior. For instance, if the weights of the MLPs ***f*** are 0, then ***f*** = **0** ⇒ *σ*(***f***) = 0.5 ·**1** ⇒ 𝒢= 2 ·*σ*(***f***) = **1**. The model can then learn more sophisticated gain patterns by diverging from the default identity gain operation. We take the square root when computing EAN-full gain map 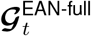 to ensure that gain values are in the same range [0, 2] as for the other models.

Dimension of 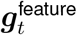 depends on the convolutional block (i.e. the number of features/channels ℝ^*c*^ in the convolutional layer). Dimension of 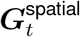 is global for all convolutional blocks 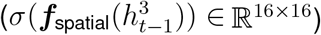, but the gain map is scaled to match the spatial dimensionality of the layer’s activation tensor using bilinear interpolation 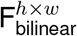.

The motivation to broadcast feature-based attention across the visual field came from canonical findings that feature-based attention produces spatially global modulation^6–9^ as well as previous modeling efforts.^10^ Similarly, the motivation to broadcast spatial attention across feature channels came from findings that spatial attention enhances neural responses at the attended location across feature preferences.^11–13^

#### Readout

On each time step *t*, we pass the 256-D hidden state of the final recurrent block 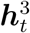 as input to the *readout* mechanism.

For the VCS task, the model makes two predictions:

- *what* prediction: linear digit classifier (256 → 10) outputting logits for each digit class
- *where* prediction: linear location predictor (256 → 100) outputting logits for positions in a 10 *×* 10 grid

Other readout heads used:

- *TinyImagenet*: linear classifier (256 → 200) outputting logits for each TinyImagenet class
- *orientation change detection* task: linear classifier (256 → 3) outputting logits for model “saccade” (“left”, “center”, “right”)
- *grating detection* task: linear classifier (256 → 2) outputting logits for binary target present judgment (“present”, “absent”)

**Full forward pass through EAN**. *t* represents the time step, while *i* represents the convolutional block index.

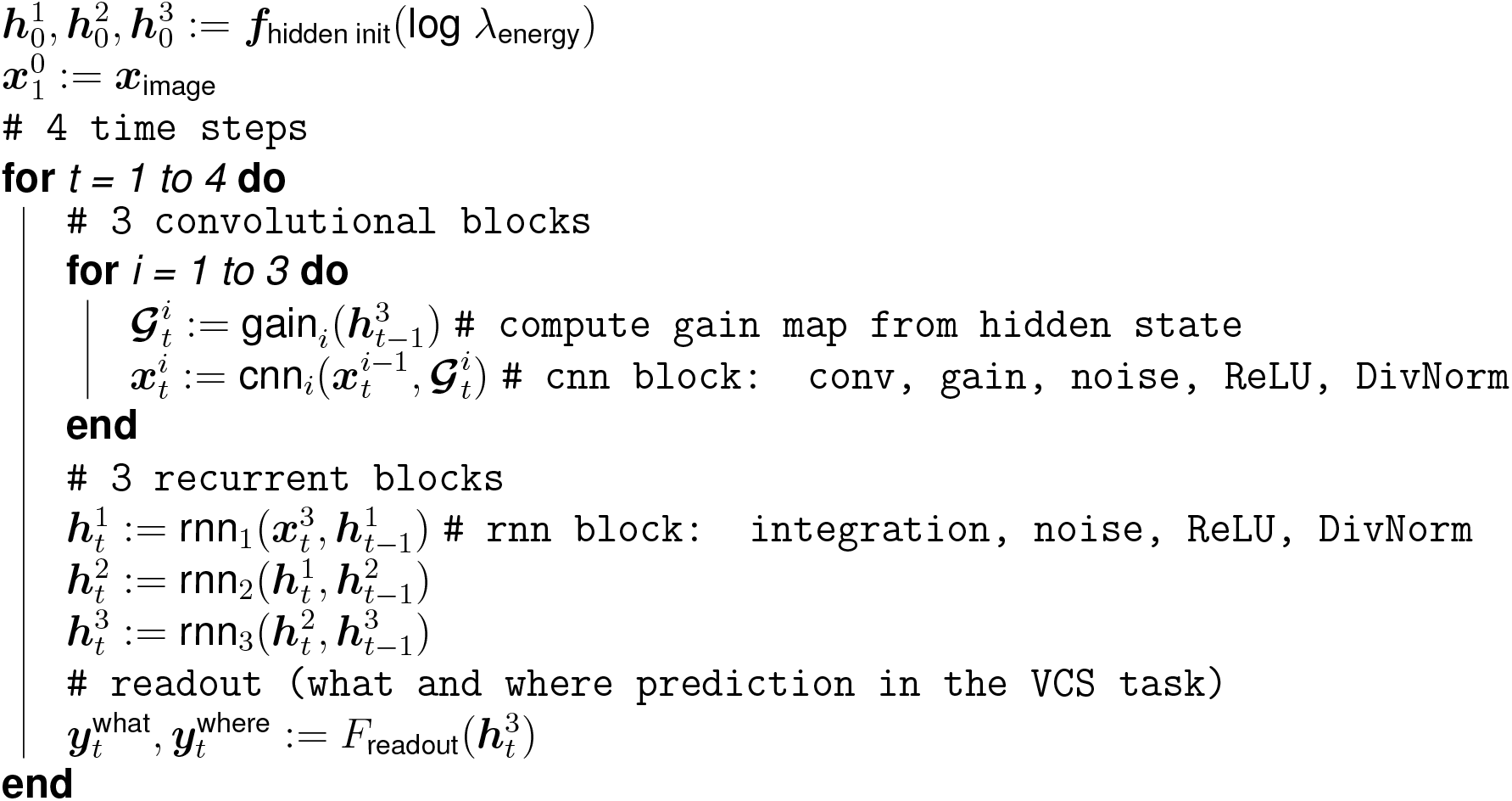

The hidden state is initialized using an MLP with 64-D hidden layer (1 → 64 → 256 × 3) that takes energy-cost factor *λ*_energy_ as input. For the first set of modeling results demonstrating that attention improves energy efficiency (Figure 4a), *λ*_energy_ is fixed for a single model instance during training: log *λ*_energy_ ∈ {−12, − 11, …, − 5}. For the second set of modeling results showing that attention enables flexible energy-accuracy trade-offs (Figure 4b), *λ*_energy_ is sampled from a distribution of energy prices on a trial-by-trial basis during training: log *λ*_energy_ ∼ Uniform(−12, −6).

#### EAN loss

Total loss for one full forward pass is the sum of losses for the individual four time steps 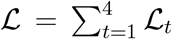. Loss for an individual time step ℒ_*t*_ includes errors for what and where components of the VCS task, energetic costs for action potentials and synaptic transmission, and a small gain regularization term:

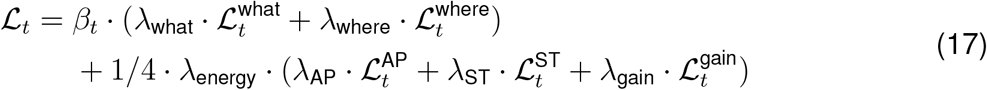

*λ*_what_, *λ*_where_, *λ*_energy_ control the trade-off between the costs of task errors and energy use. We set *λ*_what_ = 1 and *λ*_where_ = 1 since they were already in comparable ranges. *λ*_energy_ is either fixed or sampled on a trial-by-trial basis (see above). 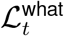 is the cross-entropy between model prediction of the digit class and one-hot label. 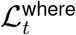 is the KL divergence between a predicted distribution over spatial locations and the “true” location map (see details in the neural network training dataset generation).

*β*_*t*_ controls the importance of providing the correct answer on a given time step. We found that giving equal weight across time steps (*β*_1_, *β*_2_, *β*_3_, *β*_4_ = 1*/*4) led to models that, to a significant degree, would not use further time steps and recurrent computation to improve their answer. In contrast, models with all the weight on the last time step (*β*_1_, *β*_2_, *β*_3_ = 0, *β*_4_ = 1) would arrive at a better answer at *t* = 4 than the constant *β*_*t*_ models, but they would not give meaningful answers for *t* = 1, 2, 3 (the model would not have any incentive to provide a correct answer for the first three time steps). To strike a balance, we set *β*_1_, *β*_2_, *β*_3_ = 0.067, *β*_4_ = 0.8 This encouraged the model to use recurrent computation to arrive at a better answer, but also incentivized the model to provide a meaningful answer for all time steps (“anytime” behavior).

For the definitions of action potential loss 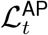 and synaptic transmission 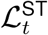, see the energy-accounting framework section below.

The gain regularization loss term 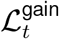 penalizes deviation of gain signals from neutral modulation (**G** = **1**). All else being equal, the regularization term nudges the gain values towards neutral ones. It also prevents degenerate EAN-full solutions where many feature–spatial gain combinations can produce identical modulation patterns (e.g., 0.8 (feature) × 1.25 (space) = 2 (feature) × 0.5 (space) = 1). For all gain types, the regularization loss term is the mean absolute deviation of the pre-scaled gain values from 0.5 (the sigmoid value corresponding to identity gain). For feature-based gain this is averaged across layers and channels:

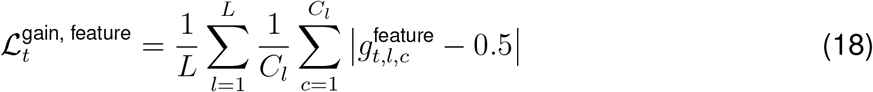

for spatial gain across spatial locations:

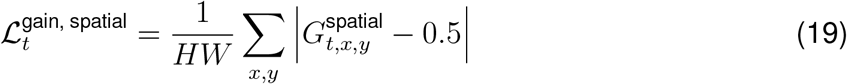

and for temporal gain applied to the scalar gain value directly:

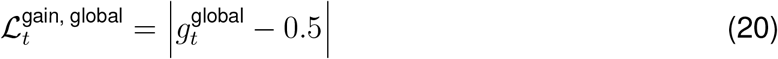

The total regularization term sums over active gain types. We set *λ*_gain_ = 1000, which results in gain regularization term 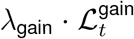 being one order of magnitude smaller than the sum of the energy costs for action potentials and synaptic transmission, acting as a gentle regularizer on the gain. Note that the gain regularization term is not included in the reported energy use of the models.

### Energy-accounting framework

The energy term consists of two components 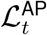 and 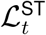, weighted by *λ*_AP_ and *λ*_ST_. The ratio between energy used for synaptic transmission and action potentials in real brains is approximately 3-to-1.^14^ To maintain this ratio in our model, we set *λ*_AP_ = 1.0 and *λ*_ST_ = 0.025, so that 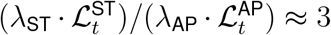 during TinyImagenet pre-training. We do not include resting potentials^14^ among the energy cost terms since all units in our models maintain the ability to produce activations—resting potential cost would be approximately constant across models.

#### Action potentials

To account for action potentials, we treat post-ReLU activations as proportional to neural firing rates^14,15^ and use their sum as the energetic cost, reflecting the total metabolic cost of generating spikes:

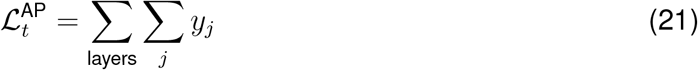

where *y*_*j*_ are the post-ReLU activations summed over all units *j* per sample, flattening whatever dimensions are present (spatial and channel dimensions for convolutional layers; hidden or output dimensions for RNN and MLP layers). This is consistent with efficient coding models and standard sparseness penalties on unit activity.^16^

#### Synaptic transmission

Previous work has used L1-norm of synaptic weights^17^ or the sum of absolute pre-activations^18^ as proxies. However, these measures have important limitations.

While an L1 penalty on the weights promotes structural sparsity, it does not account for activity-dependent transmission costs, which depend on *both* the synaptic weight and the pre-synaptic firing rate. A large weight incurs metabolic cost only when the pre-synaptic neuron is active, so it may be cheap if the pre-synaptic neuron is highly selective and thus rarely active.

Measuring pre-activations operates at the wrong level of granularity. Pre-activation represents the net input to a neuron after excitatory and inhibitory signals have already been summed. This means high synaptic transmission costs can be masked when many individual synaptic events cancel out.

To accurately capture the costs of synaptic transmission, we measure activity at the level of individual synapses *before* this summation occurs, that is, the absolute products of pre-synaptic firing rates and synaptic weights (Fig. 2d). We implement these biologically plausible energy measures for action potentials and synaptic transmission across *all* EAN components, including the convolutional layers in the visual hierarchy and recurrent layers in the attentional controller.

For a linear layer (used in MLPs and recurrent blocks) with input ***x*** ∈ ℝ^*N*^ and weights ***W*** ∈ ℝ^*N ×M*^, synaptic the synaptic transmission cost is the sum of absolute products of inputs and weights:

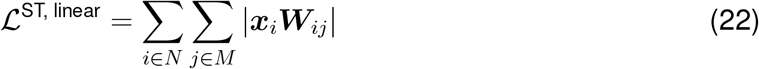

For a convolutional layer (used in convolutional blocks) with input ***x*** ∈ ℝ^*W ×H×C*^ and convolutional weights 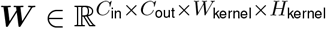:

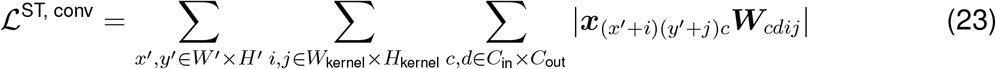

Here (*x*^*′*^, *y*^*′*^) index output spatial locations over the *W*^*′*^ *× H*^*′*^ output map, (*i, j*) index positions within the *W*_kernel_ *× H*_kernel_ convolutional kernel, and *c* ∈ *C*_in_, *d* ∈ *C*_out_ index input and output channels respectively. The input activation ***x***_(*x*_*′*_+*i*)(*y*_*′*_+*j*)*c*_ is the pre-synaptic firing rate at the spatial position in the input map corresponding to the kernel offset (*i, j*) relative to output location (*x*^*′*^, *y*^*′*^).

Computing synaptic transmission costs for every synapse would be prohibitively expensive, so we develop a stochastic estimator that samples synapses during training. For linear layers, we sample 5% of input neurons and 5% of output weights; for convolutional layers, we sample 5% of input and output channels. The estimates are unbiased: sampled sums are rescaled by the inverse sampling ratio to recover the full synaptic transmission cost in expectation. This enables gradient-based optimization of the full model while maintaining a comprehensive account of metabolic costs throughout the network. To ensure stable training, the energy costs 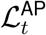 and 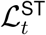 are annealed gradually during training (see section on annealing below).

#### Noise

Biological neural systems contend with multiple inherent sources of noise—ion channel noise, synaptic noise, and thermal noise^19^—that corrupt neural signals. We add normally distributed noise (*σ*_noise_ = 0.1) to the pre-activations of all model components (convolutional, recurrent, and MLP layers), simulating intracellular voltage fluctuations and serving as a proxy for the aggregate effect of many small synaptic noise events. The presence of noise establishes a biologically realistic, continuous trade-off between signal precision and energy: larger activations yield higher signal-to-noise ratios but incur greater metabolic cost, making the action potential and synaptic transmission terms in the loss meaningful throughout the network. In convolutional blocks, noise is applied *after* gain modulation, consistent with neurophysiological findings that gain modulation scales the signal while preserving internal noise levels.^20^ To ensure stable training, noise magnitude is annealed gradually during training (see section on annealing below).

#### Plotting energy use

All reported model energy use sums action potentials and synaptic transmission for all components of the model across all time steps for a single trial:

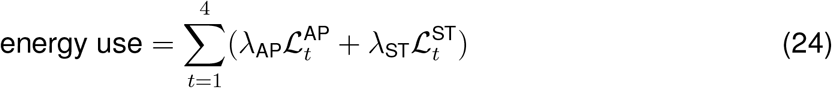

To demonstrate energy-accuracy trade-offs (Fig. 4), we fit the generalized logistic (Richards) function to log average trial energy-use against average trial what and where accuracy across the energy-factor values *λ*_energy_:

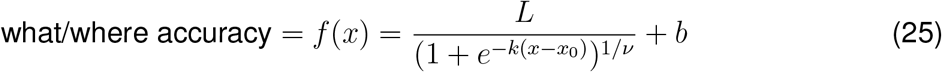

where *x* is the log average energy-use. The error bars in the energy-accuracy plots are 95% confidence intervals computed via bootstrapping (n=1000) showing variability across instances.

To plot energy savings, we inverted the logistic fits for each accuracy level and plot how much of the baseline energy is saved by each EAN version with respect to baseline.

### Training

#### Noise and energy cost annealing during training

We found that immediate introduction of full magnitude noise and energy costs during optimization destabilized optimization. We therefore “anneal” both noise and energy costs gradually, scaling each by a weight *w*_epoch_ ∈ [0, 1] that increases linearly from zero:

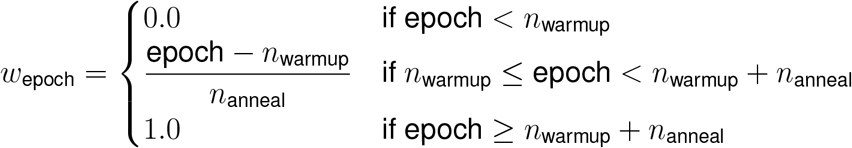

where *n*_warmup_ is the number of epochs before any noise or energy cost is applied, and *n*_anneal_ is the number of epochs over which the weight increases from 0 to 1. The same procedure is applied independently to noise and energy cost, with separate warmup and annealing epochs for each. Noise and energy costs have the opposite effect on weight magnitudes. Noise incentivizes increasing weights to maintain high signal-to-noise ratio. Energy costs incentivize decreasing weights. For all tasks and datasets (TinyImagenet pre-training, VCS, orientation change detection, grating detection task), we first anneal the noise (which increases weight magnitude) before annealing the energy cost (which decreases their magnitude).

#### TinyImageNet pre-training

Biological visual systems learn general-purpose representations that can be flexibly deployed across many tasks.^21^ We hypothesized that attentional mechanisms should adapt general visual features to specific task demands, rather than the visual hierarchy itself being optimized for a single task. Consistent with this, we found that models trained end-to-end on VCS developed filters that discriminated digits from letters already in the first convolutional layer—a biologically unrealistic degree of task specialization. We therefore pre-train the visual hierarchy on object classification using the TinyImageNet dataset^22^ (200 classes, 100k train / 10k validation images), then freeze its weights before training the attentional controller and readout on VCS. This fixes the pre-attentional selectivity of all units in the visual hierarchy, so that each downstream task adapts only the attentional controller and gain mechanisms.

Pre-training used a single feedforward time step without gain mechanisms, a 200-way softmax readout, and a small energy-cost factor (sampled during training log *λ*_energy_ ∼ Uniform(− 12, − 9)) to encourage energy efficiency of the base feedforward visual hierarchy. We converted images to grayscale and used image augmentation techniques during training to prevent overfitting: random resized cropping, horizontal flipping, TrivialAugmentWide,^23^ normalization using Tiny-Imagenet statistics, and random erasing. We trained for 300 epochs with batch size 128 using AdamW^24^ (learning rate 5 × 10^−4^, betas [0.9, 0.999], weight decay 0.0) with a StepLR scheduler (step size 150, *γ* = 0.5). We set the noise anneal parameters to *n*_warmup_ = 80, *n*_anneal_ = 5, and energy anneal parameters to *n*_warmup_ = 100, *n*_anneal_ = 10. The pre-trained model achieved ∼ 27% top-1 validation accuracy—lower than state-of-the-art, which is expected given relatively small model size, added noise, and energy regularization.

#### Training the attentional controller and gain mechanisms for VCS

After pre-training the model on Tiny ImageNet, we froze the weights in convolutional layers, and only adapted the attentional controller (including the hidden state initialization MLP), MLPs implementing gain mechanisms, and readout layers.

We trained all models for 200 epochs on our task (where each epoch consisted of 60k input images, batch size 128). For each batch, we generate novel search images using the digits and letters from MNIST and EMNIST training splits.

We used AdamW with the same hyperparameters as for pre-training. For noise annealing, we set 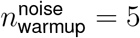 and 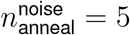. For energy cost annealing, we set 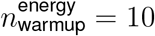 and 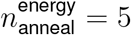. We applied StepLR rate scheduler with step size 80 and gamma 0.5.

In the *λ*_energy_-fixed regime, we fixed the energy-cost factor *λ*_energy_ for each training run:

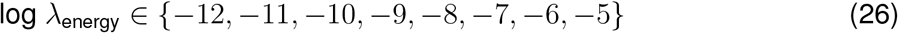

and trained 5 model instances from different random seeds per energy-cost factor per model type, resulting in 5 (model types)*×*5 (instances)*×*8 (energy cost factors) = 200 (total instances trained).

In the *λ*_energy_-flexible regime, we sample log *λ*_energy_ ∼ Uniform(− 12, − 6) during training and train 5 model instances from different random seeds per model type, where each instance is exposed to the whole distribution of energy-cost factors: 5 (model types) × 5 (instances) = 25 (total instances trained).

### Human-model VCS comparisons

#### Qualitative model behavior

For the qualitative comparison between human and model behavior (Fig. 5a), we plot model performance at a single intermediate energy cost (log *λ*_energy_ = − 7.7) from the *λ*_energy_-flexible regime. Model accuracy is plotted as a function of time step (left column, top axis) alongside human accuracy as a function of presentation time (left column, bottom axis), and as a function of number of distractors (right column). As a qualitative proxy for human difficulty judgments, we use model prediction entropy in the what dimension. Unless otherwise noted, all model accuracies reported throughout the paper are from the final time step (*t* = 4); this panel is the exception, where we show accuracy across all time steps to illustrate the dynamics of recurrent inference. Shaded regions reflect 95% bootstrapped confidence intervals across model instances and 400 search images from the VCS behavioral dataset.

#### What and where error consistency

We used Cohen’s *κ*^25^ to measure “error consistency”^26^ between pairs of classifiers (human–human or human–model), defined as:

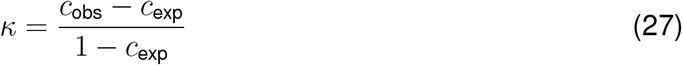

where *c*_obs_ is the observed proportion of trials on which both classifiers gave the same response (both correct or both incorrect), and *c*_exp_ = *a*_1_*a*_2_ + (1 − *a*_1_)(1 − *a*_2_) is the proportion expected by chance given their respective accuracies *a*_1_ and *a*_2_. Error consistency thus measures trial-by-trial agreement while controlling for agreement expected by chance given the marginal accuracies of each classifier.

For human–human error consistency, we computed *κ* for every pair of subjects on the trials they both completed, separately for the what and where components, and report the mean across all pairs. The shaded region in Fig. 5b for human–human error consistency represents 95% bootstrapped confidence intervals (n=10,000), reflecting variability in error consistency across different pairs of subjects.

For human–model error consistency, we used model predictions from the final time step (*t* = 4) in the *λ*_energy_-flexible regime. For each model type trained in the *λ*_energy_ flexible regime, we computed *κ* between each model instance and each human subject, for each energy-cost factor *λ*_energy_ used at evaluation. We report *κ* as a function of *λ*_energy_ (Fig. 5b), averaging across model instances and human subjects. Shaded regions reflect 95% bootstrapped confidence intervals.

#### Predicting human difficulty judgments

We tested whether model-derived measures could explain variance in human trial-level difficulty judgments beyond basic stimulus properties. For each trial, we averaged difficulty ratings (z-scored per subject) across subjects to obtain a single trial-level difficulty score. We used models trained under the *λ*_energy_-flexible regime.

We first quantified variance explained by stimulus parameters alone using ordinary least squares (OLS) regression with the number of distractors and image noise as predictors. We then tested whether model-derived covariates—trial-level energy use and prediction entropy (entropy of the what and where output distributions)—explained additional variance beyond stimulus parameters. For each model variant and energy-cost value *λ*_energy_, we fit three nested regression models: (1) stimulus parameters plus model energy use, (2) stimulus parameters plus prediction entropy, and (3) stimulus parameters plus both energy use and prediction entropy. Additional *R*^2^ was computed as the difference in *R*^2^ between each nested model and the stimulus-parameters-only baseline. All predictors were standardized before fitting.

For each model variant, we selected the energy-cost factor *λ*_energy_ that maximized the additional *R*^2^ for each set of covariates (see Extended Data Fig 2 for results across the full range of energy costs).

Statistical significance was assessed using a bootstrap procedure (1,000 iterations, resampling 3,000 trials with replacement from the 389 behavioral trials per iteration). For each bootstrap sample, we fit all regression models and computed additional *R*^2^ values. To test whether each EAN variant explained significantly more variance than the baseline model, we computed the bootstrap distribution of pairwise differences in additional *R*^2^ and derived two-tailed p-values, corrected for multiple comparisons using the Bonferroni method (4 comparisons against baseline per metric).

### Electrophysiology replication

#### Cohen & Maunsell [27]: firing rate, mean-matched Fano factor, noise correlation

##### Orientation change detection task

We implemented a simplified four-time-step version of the orientation change detection task from Cohen & Maunsell [27]. On each trial, the model receives a 64 × 64 grayscale image containing two sinusoidal gratings (4 cycles, radius 8 pixels) presented simultaneously on the left (centered at pixel [25, 16]) and right (centered at pixel [25, 48]) sides of the display, viewed through Gaussian apertures (*σ* = *r/*2, where *r* = 8 is the aperture radius in pixels), forming Gabor patches. On the first time step (cue frame), an attentional cue (a small white or black 3 × 3 square at the image center) indicates which side is likely to change, and both gratings are visible left and right of the cue. On one of time steps 2–4 (uniformly sampled), the orientation of one grating changes by a random amount sampled uniformly from [1°, 90°] (with uniformly sampled sign). The change persists for the remainder of the trial. On 80% of trials the change occurs on the cued side (valid trials); on the remaining 20% it occurs on the uncued side (invalid trials).

Instead of the dual *what* and *where* readout used in VCS, we implemented a 3-way “saccade” readout (“left”, “center”, “right”). The model was trained using the cross-entropy loss to output “center” on all time steps except the change time step, and “left” or “right” on the change time step depending on which side changed.

##### Training details

As for VCS, we kept the pre-trained visual hierarchy frozen and trained only the attentional controller (EAN-full), gain mechanisms, and saccade readout under the *λ*_energy_-flexible regime. We trained for 50 epochs using AdamW (learning rate 5 *×* 10^−4^) with noise annealing 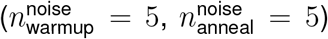 and energy cost annealing 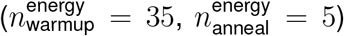.

Each epoch consisted of 2,000 trials. In the gain mechanisms, we multiplied the input to the sigmoid by a factor of 0.1, essentially reducing “gain sensitivity” to encourage more stable training. Below, we analyze the model under *λ*_energy_= −10.

To encourage the model to develop a unimodal spatial attention pattern (attending to the cued side), we initialized the cue validity probability at 100% and only decreased it to 80% for the last epoch. The cue validity schedule was designed to address a practical training challenge: the model lacked a built-in unimodal spatial attention prior. Starting with 100% valid cues allowed the model to first learn to attend to the cued side, before introducing invalid trials that required the model to occasionally detect changes at the uncued location.

##### Experimental days replication

To replicate the experimental design of Cohen & Maunsell [27], for our analysis of the trained model we fixed the pre-change orientations of both gratings within each simulated recording “day” (50 days total, each with a unique pair of orientations), with many repetitions per day (20 repetitions).

##### Activation to spike rate conversion

To compare model activations with electrophysiological recordings, we converted post-ReLU activations from convolutional layer conv3 (256 channels, corresponding to an intermediate-to-high visual area analogous to V4) to simulated spike rates. We extracted activations with receptive fields corresponding to the left and right grating centers, yielding 256 “units” per side. Activations were linearly scaled by a factor of 300 and offset by a baseline rate of 1 spike/s:

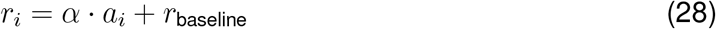

where *a*_*i*_ is the post-ReLU activation of unit *i, α* = 300 is the scaling factor, and *r*_baseline_ = 1. We then sampled spike counts from a Poisson distribution with rate *r*_*i*_ to introduce realistic trial-to-trial variability. Following Cohen & Maunsell [27], all analyses used only correctly completed trials, and activations were taken from the time step immediately preceding the orientation change (excluding the first time step, as the model has not yet processed the attentional cue).

##### Mean-matched Fano factor

We computed the mean-matched Fano factor following the procedures described in Cohen & Maunsell [27] and Churchland *et al*. [28]. For each simulated recording day, we computed the mean spike count and variance for each unit under attended and unattended conditions. We then found the common distribution of mean firing rates across all day–attention–side combinations by binning mean firing rates into 10 percentile-based bins and taking the minimum count per bin across all conditions. Units were subsampled from each condition to match this common distribution, ensuring that any differences in Fano factor between attended and unattended conditions could not be attributed to differences in mean firing rate. The Fano factor was estimated as the slope of a linear regression of spike count variance on spike count mean across the matched population. To obtain realistic Fano factor values, we divided the raw spike count variance by a factor of 20, compensating for the inflated trial-to-trial variability introduced by the activation-to-spike-rate scaling.

##### Noise correlation

Noise correlations were computed as Pearson correlations between pairs of units’ spike counts across trials, separately for each day and attentional condition. We computed within-”hemisphere” correlations (pairs of units with receptive fields on the same side) and across-”hemisphere” correlations (pairs with receptive fields on opposite sides), further split by whether the units’ receptive fields were in the attended or unattended side. For each unit, we averaged its pairwise noise correlations with all other units in the same condition (within-side attended, within-side unattended, or opposite-sides). These per-unit mean noise correlations were plotted as a function of the unit’s mean firing rate, binned into 10 equally spaced bins from 1 to 40 spikes/s. Units with firing rates exceeding 100 spikes/s were excluded.

#### Debes & Dragoi [29]: V4→V1 optogenetic feedback suppression

##### Grating detection task

We implemented a simplified four-time-step version of the grating detection task from Debes & Dragoi [29]. On each trial, the model receives a 64 × 64 grayscale image that can contain sinusoidal gratings (3 cycles, radius 5 pixels) on the left (centered at [25, 16]) and/or right (centered at [25, 48]) sides, viewed through hard-edged circular apertures (radius 5 pixels). On the first time step, an attentional cue (a small white or black 3 × 3 square) indicates which side to report. On either the second or third time step (uniformly sampled), the stimulus frame is presented; the remaining time steps show an empty gray display. The presence of a grating on each side is independently sampled (50% probability), and when present, its contrast is uniformly sampled from [0, 0.5] and its orientation is uniformly sampled from [0, *π*). Following the original experiment, the cue is 100% valid: at the end of the trial, the model must report whether a grating is present on the cued side, ignoring the other side. The model uses a binary readout (“present” vs. “absent”) and the model is trained using the cross-entropy loss.

##### Training details

We used EAN-spatial for this task and trained for 30 epochs using AdamW (learning rate 5 *×* 10^−4^) with noise annealing 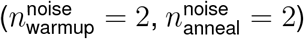 and energy cost annealing 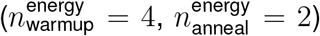. Each epoch consisted of 10,000 trials. As for the other tasks, we kept the pre-trained visual hierarchy frozen and trained only the attentional controller, gain mechanisms, and readout under the *λ*_energy_-flexible regime. We used log *λ*_energy_= − 7 for the analysis below.

##### Model “optogenetics”: feedback suppression

To model the optogenetic suppression of V4 → V1 feedback from Debes & Dragoi [29], we implemented a gain suppression procedure that stochastically dampens the top-down gain signal from the attentional controller to the visual hierarchy (model “optogenetics”). On each trial, we sample whether gain suppression is applied (50% probability, matching the laser-on/laser-off design of the original experiment). When gain suppression is active, the gain signal at the stimulus onset time step is dampened toward neutral modulation:

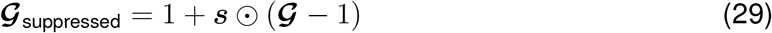

where ***s*** = exp ***z*** *γ*, ***z*** (**0, *I***) is sampled independently per unit, and *γ* = 50.0 controls suppression strength. Suppression is applied on 50% of trials at the stimulus onset time step; on the remaining trials *γ* = 0, so ***s*** = **1** and the gain is unmodified. When ***s*** = **0**, the gain is fully suppressed to neutral (𝒢_suppressed_ = 1, equivalent to removing all top-down modulation). Gain suppression was applied to cnn1 (the earliest convolutional layer, analogous to V1), consistent with the V4→V1 feedback pathway targeted in the original optogenetic experiment.

##### Analysis

We extracted activations from cnn1 (64 channels) at the spatial locations corresponding to the left and right grating centers, and converted them to firing rates using the same linear scaling procedure as for the orientation change detection task. Behavioral “target reports (%)” measures the proportion of trials the model reports that the grating is “present” at the cued side as a function of grating contrast for each combination of attentional condition (control vs. gain-suppressed). We replicated the key analyses from Debes & Dragoi [29]: (1) the effect of feedback suppression on behavioral target reports as a function of contrast, (2) the effect of feedback suppression on cnn1 firing rates (analogous to V1) as a function of time step and attentional condition, and (3) the contrast-dependence of feedback suppression effects on firing rates.

## Data and Code Availability

Data and code is available at https://github.com/eivinasbutkus/how-attention-saves-energy-in-vision.

## Acknowledgements

We thank Ilker Yildirim, Mario Belledonne, Peiyu Chen, Jackie Gottlieb, Mike Woodford, Chris Baldassano, Ruth Rozenholtz, Xue-Xin Wei, Xaq Pitkow, Alan Stocker, members of the “Cognitive and Neural Computation Lab” at Yale University, and participants of the “Multi-resource-cost Optimization of Neural Networks” working group at the NSF AI Institute for Artificial and Natural Intelligence for helpful comments and discussions on earlier versions of this work.

This work is supported by the funds provided by the National Science Foundation and by DoD OUSD (R&E) under Cooperative Agreement DBI-2229929 (The NSF AI Institute for Artificial and Natural Intelligence).

## Author Contributions

E.B. and N.K. conceived the project. E.B. developed the model, energy-accounting framework, and code. Z.Y. helped implement the TinyImagenet pre-training and provided helpful comments on early versions of the model. E.B. designed and conducted the behavioral experiment. E.B. performed all analyses. E.B. created the figures. E.B. and N.K. wrote the manuscript with feedback from Z.Y. N.K. supervised the project.

## Extended Data

**Extended Data Figure 1:**
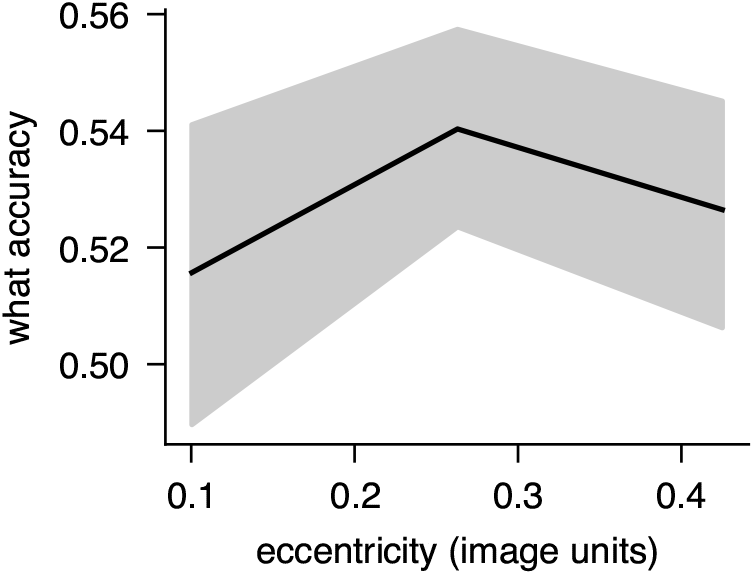
Human what accuracy does not meaningfully change as a function of target eccentricity.

**Extended Data Figure 2:**
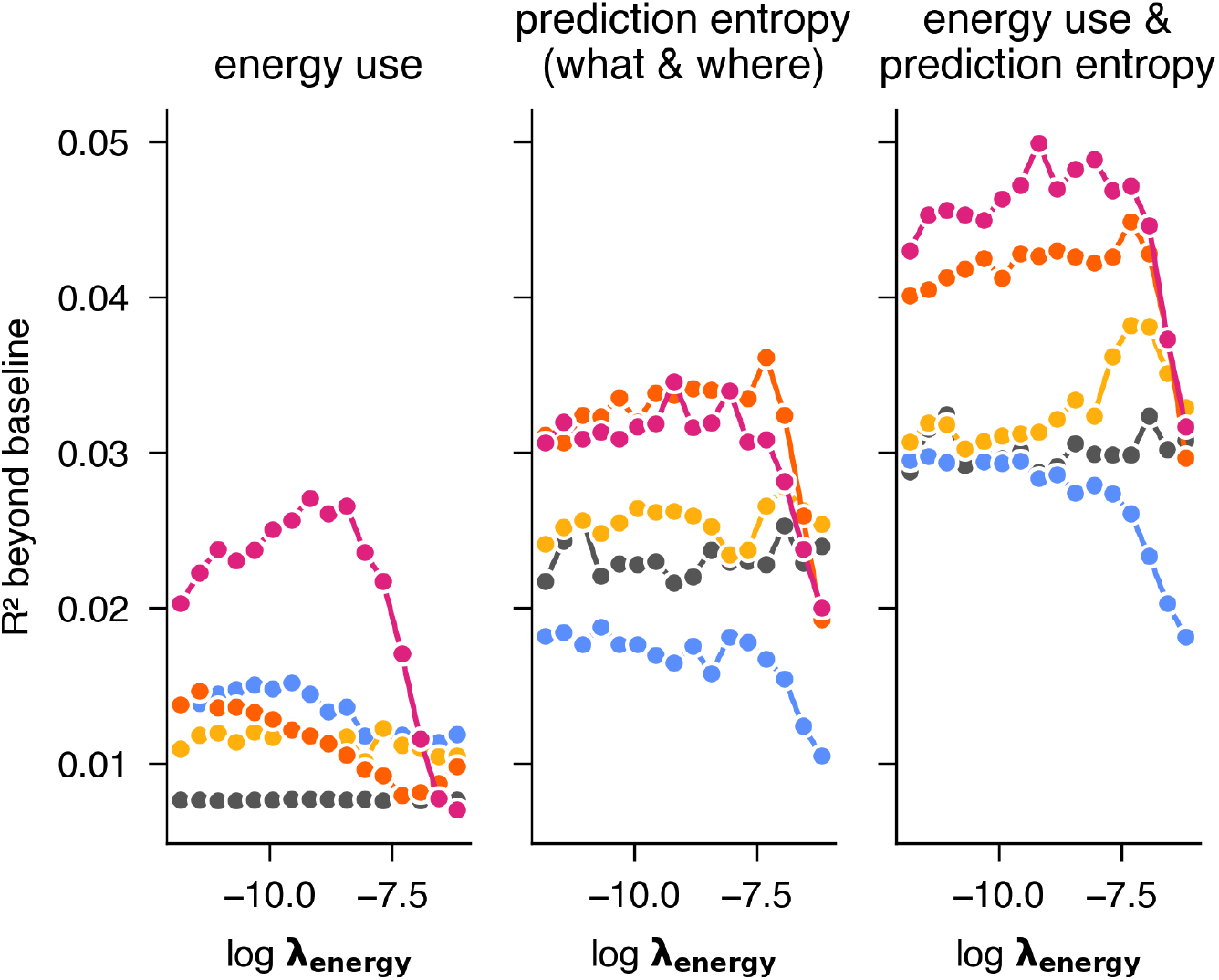
Additional *R*^2^ as a function of log *λ*_energy_. For the main results (Fig. 5c), we select *λ*_energy_ parameters that maximize additional *R*^2^.

## Supplementary Information

### Robustness of energy savings to gain application costs

Our energy-accounting framework measures the cost of *computing* gain signals through the attentional controller and gain MLPs—including synaptic transmission and action potentials in all layers of these components. However, the biophysical cost of *applying* multiplicative gain to target neurons (e.g., via dendritic or synaptic mechanisms) is not explicitly measured as a separate term.

Part of the application cost is implicitly captured: the second layer of each gain MLP produces the gain values, and its synaptic transmission cost scales with the magnitude of the gain signal. Nonetheless, the true biological cost of gain modulation at the target site may exceed what our framework currently accounts for.

To assess whether our conclusions are robust to this underestimation, we computed an upper bound on how much more expensive the non-CNN components (attentional controller and gain mechanisms) could be while still yielding net energy savings. Comparing baseline (log *λ*_energy_ = − 9, mean what accuracy = 44.7%) and EAN-full (log *λ*_energy_ = − 7, mean what accuracy = 53.8%) in the fixed *λ*_energy_ regime (Fig. 4d), the CNN visual hierarchy accounts for ∼ 96% of total energy use, while the RNN and gain mechanisms together account for only ∼ 4%. The net energy savings from attention are approximately 19 times larger than EAN-full’s total non-CNN energy expenditure. This means that even if the costs associated with the attentional controller and gain mechanisms—including any unmeasured gain application costs—were ∼ 19 × higher than currently estimated, attention would still break even energetically. Any smaller costs would preserve net savings. Note that this is a conservative bound, since EAN-full also achieves substantially higher accuracy than the baseline in this comparison.

Our energy-accounting framework is readily extensible: as empirical estimates of gain application costs become available, they can be incorporated as additional terms in the objective without modifying the overall architecture or training procedure.

